# Biodegradable Intra-arterial Devices for Focal Drug Delivery to Targeted Organs

**DOI:** 10.64898/2026.02.23.707478

**Authors:** Manas Kinra, Ruoyu Sheng, Yiqing Chen, Amancio Jose de Souza, Anil Bhatia, Garrett Sakomizu, Jinrui Tan, Dongwei Sun, Edward Zagha, Huinan Liu

## Abstract

This study presents the development of biodegradable intra-arterial drug delivery (IADD) devices for the focal treatment of targeted organs. The IADD devices are fabricated using magnesium (Mg) and poly(glycerol sebacate) (PGS), leveraging their biocompatibility and tunable biodegradability, and are loaded with two model drugs, dexamethasone (DEX) or cisplatin (CIS). The IADD devices with helical and linear designs were fabricated for focal drug delivery to targeted organs and characterized for their microstructure and composition using scanning electron microscopy (SEM), energy-dispersive X-ray spectroscopy (EDS), thermogravimetric analysis (TGA), and Fourier-transform infrared spectroscopy (FTIR). The results confirmed the successful incorporation and stability of the drugs within the device. The IADD devices demonstrated in vitro release of DEX and CIS over 30 days, with drug-dependent and fabrication-dependent release profiles. The IADD devices demonstrated cytocompatibility with endothelial cells and sustained pharmacological activity against glioma cells throughout the *in vitro* release period. DEX-loaded IADD devices were implanted into the artery upstream of a target organ in rat models. The devices implanted into the renal artery to target the kidney and the carotid artery to target the brain achieved 109-fold and 68-fold improvements, respectively, in organ vs systemic drug levels compared to oral drug administration. Arterial histology and explanted-device analysis provided preliminary evidence of local vascular tolerability and device integrity over the examined implantation period. Overall, these results support IADD devices as a proof-of-concept approach for focal intra-arterial drug delivery to targeted organs, demonstrating substantially elevated target-organ drug levels relative to systemic exposure.

**Highlights:**

- Introduction of a novel intra-arterial drug delivery (IADD) platform
- IADD devices were fabricated from bioresorbable Mg and drug-loaded PGS
- DEX- and CIS-loaded IADD devices were evaluated for in vitro release and bioactivity
- IADD delivery achieved marked DEX enrichment in the kidney and brain
- Post-implantation studies support preliminary vascular safety

## 1. Introduction

The development of focal drug delivery systems represents a significant advancement in treating diseases localized to specific tissues or organs. Compared with conventional methods, such as intravenous (IV) and oral administration, these systems allow for precise medication delivery directly to the targeted area while reducing systemic exposure and off-target side effects [1]. Several delivery techniques have been proven to achieve focal drug delivery, including intra-arterial (IA) injectables [2], pumps [3], and implants [4]. Studies show that IA chemotherapy achieves markedly higher local drug concentrations while reducing systemic exposure compared to conventional systemic routes. For example, IA cisplatin infusion in high-grade glioma patients yielded 8-12-fold higher tumor platinum levels and ∼30% lower systemic levels (i.e., area under the curve, AUC) than intravenous dosing [5, 6]. Likewise, hepatic artery infusion of oxaliplatin or albumin-bound paclitaxel achieved 5-10-fold higher tumor/plasma ratios with 15-60% lower systemic exposure [7, 8]. In a recent single-center series of 70 glioblastoma patients undergoing 139 super-selective IA cerebral infusions, the completion rate of intended treatments was 95.7% and drug-related adverse events occurred in only 8.6% of cases [9]. Collectively, these findings underscore the pharmacokinetic and safety advantages of focal IA delivery of maximizing on-target effect while minimizing off-target exposure. However, current IA technology allows for acute delivery only, limited to the duration of endovascular catheterization.

The objective of this work is to develop and evaluate proof-of-concept biodegradable intra-arterial devices for focal drug delivery. In this study, we designed novel biodegradable intra-arterial drug delivery (IADD) devices, which could be placed into large- or middle-sized arteries via minimally invasive endovascular procedures and provide tunable release profiles for focal drug delivery to any organ perfused by a large- or middle-sized upstream artery. Compared with direct organ implants, IADD devices can be implanted endovascularly, offering a less invasive option while still achieving high local drug concentrations in the targeted downstream organs. Moreover, our IADD devices can be loaded with a wide range of therapeutics and fabricated into a variety of geometries such as helical structures, linear plank structures, half to full ring structures, or stent-like structures for controlling the amount of drug loading and release. Additionally, IADD devices have the potential for broad organ perfusion, compared to the diffusion-limited dispersal of direct organ implants [10].

For the first-generation prototypes of our novel IADD device, we selected magnesium (Mg) as the base material and poly(glycerol sebacate) (PGS) as the drug-loading polymer for the following reasons. Both Mg and PGS are biodegradable materials [11–13] eliminating the need for a subsequent procedure to remove the device, thereby minimizing potential damage to the implanted vessel [14]. Mg serves as the backbone of the device, providing the mechanical stiffness and strength required for endovascular delivery and for retaining the shape of the device in the artery. Mg is widely used in biomedical implants due to its biocompatibility and biodegradability, as well as mechanical properties [15–17] . Furthermore, Mg itself may improve intra-arterial drug delivery, as previous studies of Mg in vascular stents have shown that the released Mg^2+^ enhances blood flow, inhibits platelet activation, and prevents vasoconstriction [18]. Mg is fully bioresorbable, excreted through urine and feces as Mg^2+^ ions [19, 20]. PGS has gained prominence in tissue engineering and drug delivery due to its customizable and straightforward synthesis process [21]. PGS degrades via surface erosion in the body fluids, which affords stable drug release compared with polymers that degrade via bulk erosion such as poly-lactic acid (PLA) and poly-glycolic acid (PGA) [22]. The degradation products, glycerol and sebacic acid, are biocompatible and are naturally found and metabolized in the human body [23]. The synthesis conditions of PGS [24], such as reaction times and temperatures, can be fine-tuned to control the degree of esterification, allowing for precise tailoring of the chemical and mechanical properties of the synthesized polymer as well as its degradation behavior [25]. Recent PGS studies further emphasize that synthesis route, curing conditions, and polymer modification can substantially influence PGS structure, degradation behavior, and drug-release kinetics, supporting the ability to optimize polymer processing parameters for each device-drug formulation [26–28]. Notably, PGS is nonimmunogenic and has shown to be non-cytotoxic, causing minimal inflammatory response [29]. In addition, the mechanical properties of PGS, which are similar to blood vessels, can minimize the mechanical mismatch between the IADD device and arteries [13]. These characteristics make our IADD devices conducive for sustained intra-arterial drug delivery.

To demonstrate the potential of our novel IADD devices, we selected two model drugs—dexamethasone (DEX, C_22_H_29_FO_5_) and cisplatin (CIS, Pt (NH_3_)_2_Cl_2_)). Both DEX and CIS are acknowledged not only for their effectiveness, but also for their high systemic toxicity. DEX, a synthetic corticosteroid, is used to treat inflammatory and autoimmune diseases [30], allergic reactions [31], respiratory issues [32], endocrine disorders, and certain cancers [33]. However, its use is restricted due to severe side effects including metabolic disorders such as obesity and insulin resistance, hypertension, and susceptibility to infections [34]. CIS, a chemotherapeutic agent, is widely used against solid tumors by binding to DNA and inducing apoptosis, but its clinical application is limited by nephrotoxicity and other severe toxicities [35]. By delivering these drugs intra-arterially to the target organs, IADD devices could enhance target organ drug levels and reduce systemic exposure, thereby increasing drug efficacy and minimizing harmful off-target side effects.

To evaluate this approach, we fabricated and characterized DEX-loaded and CIS-loaded IADD devices with two prototype geometries. We assessed device microstructure, drug incorporation, in vitro release, endothelial cytocompatibility, and released-drug bioactivity. We then tested DEX-loaded IADD devices in vivo by implanting devices in the renal artery or carotid artery of rats to evaluate focal drug biodistribution relative to oral DEX administration. Kidney and brain were selected as representative target organs because they differ substantially in vascular permeability and tissue barriers. The current study is a proof-of-concept evaluation of IADD-mediated focal drug delivery and target-organ enrichment and provides a foundation for future optimization of drug loading, release kinetics, dose-normalized pharmacokinetics, vascular safety, and efficacy testing in disease-relevant models.

## 2. Methods

### 2.1 Fabrication of Drug-loaded IADD Devices and Controls

Two structural variants of intra-arterial drug delivery (IADD) devices (helical or linear shaped structure) were fabricated using magnesium (Mg) wire cores and poly(glycerol sebacate) (PGS) coatings (Supplementary Figure S1). Full fabrication parameters including polymer synthesis conditions, curing durations, device dimensions, and sterilization are provided in the Supplementary Methods. Briefly, PGS pre-polymer was synthesized from sebacic acid and glycerol following a modified polycondensation procedure [21]. The clean Mg wires of two model geometries (helical or linear shape) were embedded within the pre-PGS matrix and vacuum-cured at 120 °C to produce the IADD devices.

Two distinct drug-loading methods were employed: a two-step post-soaking method, in which pre-cured IADD devices were sequentially soaked in ethanol and drug solution (DEX in ethanol, CIS in DMF), and a one-step co-curing method, in which 10 % w/w of DEX was directly dispersed in the pre-PGS before curing. The helical devices were first developed using the two-step method to implant into renal arteries for focal drug delivery to kidney in rats (kidney cohort 1), but their extremely high amount of drug loading and release motivated the design of smaller, linear plank devices using both methods above. The linear devices fabricated by the one-step method were also implanted into renal arteries for focal drug delivery to kidney in rats (kidney cohort 2). The linear devices fabricated by the two-step method were used for physicochemical characterization, in vitro drug release, and in vivo drug delivery to the brain in rats. Because geometry, loading method, and implantation site were not varied independently, comparisons across these cohorts were interpreted as prototype-specific outcomes rather than as a factorial comparison of individual design variables.

### 2.2 Characterize Microstructure and Composition of Drug-loaded IADD Devices and Controls

The morphology of the devices was first examined using a 3D laser scanning microscope (Keyence VK-X150). Scanning electron microscopy (SEM) was then used to analyze the surface microstructure and the cross-sections of the drug-loaded IADD devices and non-drug-loaded controls. The devices were sputter-coated with a conductive layer of gold at 20 mA for 30 s (Sputter Model 108, Cressington Scientific Instruments Ltd., Watford, UK) before SEM. The cross-section morphology was examined using a Nova NanoSEM 450 (FEI Co., Hillsboro, OR, USA) equipped with an X-Max50 detector and AZtecEnergy software (Oxford Instruments, Abingdon, Oxfordshire, UK). Energy-dispersive X-ray spectroscopy (EDS) was used to analyze the elemental compositions of the drug-loaded IADD devices and controls. An accelerating voltage of 15 kV was used for SEM imaging and EDS analysis.

### 2.3 Thermogravimetric Analysis (TGA) of Drug-loaded IADD Devices and Controls

Thermogravimetric analysis (TGA) was conducted using a TG 209 F1 Libra instrument (Netzsch). First, the pre-cured PGS blocks with a dimension of 10 mm x 10 mm x 0.6 mm were cut out and loaded with DEX or CIS following the same two-step method as described above for the IADD device fabrication. Next, 10 mg of each sample was cut out from the block and loaded into alumina crucibles and heated from 200°C to 800°C at a heating rate of 10°C/min in an air flow rate of 20 mL/min for TGA. The TGA of DEX and CIS were performed in parallel for comparison.

### 2.4 Analyze Chemical Bonding of Drug-loaded IADD Devices and Controls

Fourier-transform infrared spectroscopy–attenuated total reflectance (FTIR-ATR) was utilized to analyze the chemical bonding of drug-loaded IADD devices, IADD control device without drugs, and drug-only controls. ThermoScientific Nicolet iS10 FTIR instrument was used to measure the transmittance of the samples and controls with the wavenumbers ranging from 4000 cm^-1^ to 400 cm^-1^. The drug-loaded PGS layer of 10 mg was scraped from drug-loaded IADD devices using a blade. Similarly, a 10 mg PGS layer without drug was scraped from the IADD control devices. DEX and CIS powders were also measured as drug-only controls. The FTIR-ATR spectra of the drug-loaded IADD devices were compared with the spectra of IADD controls and drug-only controls to determine chemical bonding and drug loading.

### 2.5 Measure Drug Release Profiles and Degradation of Drug-loaded IADD Devices and Controls *In Vitro*

The drug release profiles and degradation of the drug-loaded IADD devices were analyzed *in vitro* using the immersion method. Specifically, the linear drug-loaded IADD devices (n=5) fabricated by the two-step method and controls were placed in individual wells of a 48-well plate and immersed in physiologically relevant solutions. The DEX-loaded IADD devices were immersed in revised-simulated body fluid (rSBF) and the CIS-loaded IADD devices were immersed in artificial cerebrospinal fluid (aCSF). The rSBF and aCSF solutions were prepared according to the respective composition reported in literature [13, 36, 37]. The plates were placed inside a rotary shaker (Benchmark Incu-shaker Mini) at 120 rpm and incubated at 37 °C. The supernatants were collected from each well by complete media replacement (1 mL) every 24 h for 30 days.

To further estimate recoverable DEX payload from the one-step loading method, an additional 60-day in vitro release study was performed using one-step DEX-loaded IADD devices fabricated with 10% w/w DEX relative to device mass. Devices were incubated under the same general release conditions, and release medium was collected at defined intervals through day 60. DEX was quantifiable in the release medium through day 50 and was undetectable at later time points. Therefore, cumulative DEX recovered through day 50 was compared with the DEX mass introduced during fabrication to estimate the fraction of recoverable DEX payload. This calculation was used as a recovery-based estimate of DEX loading for the one-step formulation. Because DEX became undetectable in the release medium after day 50, the cumulative DEX recovered through day 50 was interpreted as the measurable released payload under these in vitro conditions. However, this approach does not directly define spatial drug distribution within the cured PGS matrix and does not replace future validation using direct extraction.

To measure the DEX and CIS released into the media, liquid chromatography tandem mass spectrometry (LC-MS/MS) was used. DEX and CIS concentrations in daily release media were quantified using LC-MS/MS. For DEX, samples were extracted with ethyl acetate containing dexamethasone-d5 as an internal standard, dried, and reconstituted in methanol before analysis on a Waters I-class UPLC coupled to a Synapt G2-Si Q-TOF MS operated in ESI+ mode. CIS quantification followed a derivatization-based method (Shaik et al., 2017): samples were reacted with palladium acetate and diethyldithiocarbamate (DTCC), extracted with acetonitrile containing 8-Cyclopentyl-1,3-dipropylxanthine (DPCPX) as the internal standard, dried, and reconstituted prior to analysis on a Waters H-class UPLC-G2-XS Q-TOF MS in ESI+ mode. Both assays used standard curves for absolute quantification and lock-spray mass correction to ensure mass accuracy. Full extraction protocols, LC gradients, MS parameters, and MS/MS transitions are provided in the Supplementary Methods.

Mg^2+^ ion concentrations in the collected media were measured as a direct indicator of the IADD device degradation. Inductively coupled plasma optical emission spectrometry (ICP-OES; Optima 8000, PerkinElmer, Waltham, MA) was used to measure Mg^2+^ ion concentrations. First, 30 µL of media collected from each sample was diluted in deionized water (Millipore) to 3 mL. The diluted solutions were then fed into the ICP-OES using an autosampler. Mg^2+^ ion concentrations were calculated using the ICP-OES results multiplied by the dilution factor (100x).

### 2.6 Determine Cytocompatibility of Drug-loaded IADD Devices and Controls with HUVEC Cells *In Vitro*

Human umbilical vein endothelial cells (HUVEC) were used as the model cell to test the cytocompatibility of the drug-loaded IADD devices and controls in vitro. HUVECs were maintained in Endothelial Cell Growth Medium (Cell Applications Inc.) with 1% Penicillin-Streptomycin (P/S, Corning) solution. The cells were maintained in a standard incubator at 37°C with 5% CO_2_.

The DEX-loaded IADD devices, CIS-loaded IADD devices, and non-drug-loaded control devices were prepared using the two-step method as described in Section 2.1 and disinfected prior to *in vitro* cell study. Cytocompatibility of the devices was studied using the exposure culture method [38]. Specifically, the cells were first seeded into a 12-well plate at a density of 8,000 cells/cm² and cultured for 24 h. Afterward, the cells were exposed to the drug-loaded IADD devices (n=3), ethanol-IADD control devices (n=3), and DMF-IADD control devices (n=3) for another 24 h. Each device of interest was placed in a Transwell insert (Corning) and introduced to each well. The control wells (n=3) were the cells only exposed to the inserts without any devices. After exposure culture, Sulforhodamine B (SRB) assay was used to assess cell viability. The cells in each well were fixed with 1 mL of ice-cold trichloroacetic acid (10% w/v) and incubated at 4 °C for 1 h. The culture wells were washed with water and air-dried. SRB solution (0.057% w/v, 2 mL) was added to each well and incubated for 30 min, followed by washing with 1% v/v acetic acid. The wells were dried, and 3 mL of 10 mM TRIS base (pH 10.5) was added to each well. Absorbance was measured at 510 nm using a microplate reader [39]. The percentage of cell viability was calculated as follows:

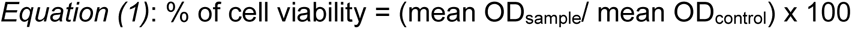

The mean OD_sample_ is the mean absorbance values of the triplicates samples; the mean OD_control_ is the mean absorbance values obtained from triplicate control measurements.

### 2.7 Evaluate Chemotherapeutic Potential of CIS-loaded IADD Devices and Controls *In Vitro*

F98 glioma cells (CRL-2397, ATCC) were used to assess the anti-cancer activity of CIS released from the CIS-loaded IADD devices using SRB assay [39]. F98 rat glioma cells were cultured in Dulbecco’s Modified Eagle Medium (DMEM, Sigma-Aldrich) with 10% Fetal Bovine Serum (FBS, Gibco) and 1% Penicillin-Streptomycin (P/S, Corning) solution. The cells were seeded into a 96-well plate at a density of 5,000 cells/cm^2^ and cultured in 150 μL of media. After 24h, 50 µL of CIS-containing aCSF solution collected from the drug release study (as described in section 2.5) was added to each well. The immersion solution collected from the non-drug-loaded IADD devices were used as controls. After 24 h, the media from each well was collected, and 30 µL of media from each sample was diluted (1:100) with deionized water to 3 mL for ICP-OES analysis of Mg^2+^ and Ca^2+^ ion concentrations. The cells were stained using the same method described in section 2.7. The experiment was performed in triplicate, and results are reported as Mean ± SD. The percentage of cell viability was calculated the same as shown in Equation 1.

### 2.8 Assess Focal Delivery of DEX to the Targeted Kidney or Brain *in Vivo* using Rat Models

Male Sprague Dawley rats (180-200 g) and male Fischer rats (180-200 g) procured from Charles River Labs were used to assess the focal drug delivery to the kidney and brain, respectively. All experimental procedures were approved by the Institutional Animal Care and Use Committee (IACUC) of the University of California, Riverside. Animals were housed in groups of three under controlled conditions with a 12 h light/dark cycle and free access to food and water.

#### 2.8.1 Focal delivery of DEX to the kidney in rats

Sprague Dawley rats were divided into two experimental cohorts. Both cohorts were used to evaluate focal drug delivery of DEX from the IADD devices to kidney versus oral delivery. Cohort 1 consisted of the rats that were implanted with helical-shaped DEX-loaded IADD devices (fabricated by the two-step method) in the left renal artery (n=2) and the corresponding rats that received DEX (1 mg/mL) in drinking water as oral controls (n=3). Cohort 2 consisted of the rats that were implanted with linear-shaped DEX-loaded IADD devices (fabricated by the one-step method) in the left renal artery (n=3) and their oral control group (n=3).

All surgical instruments were sterilized using an autoclave at 121°C and 15 psi for 30 min and handled aseptically thereafter. Animals were weighed before the procedure, and pre-operative analgesics, buprenorphine extended release (0.65 mg/kg s.c.) and meloxicam (2 mg/kg s.c.), were administered 30 min prior to anesthesia. Rats were anesthetized with 4% isoflurane in 1 L/min O_2_, followed by ketamine (80 mg/kg i.p.) and xylazine (10 mg/kg i.p.). They were then transferred to a controlled heating pad with isoflurane maintained at 1% in 1 L/min O₂, and veterinary ophthalmic lubricant was applied to their eyes. An i.p. injection of 5 mL/kg normal saline was given pre-operatively. Animals were positioned supine with limbs fixed using low-tack adhesive. After confirming adequate anesthesia by monitoring reflexes, abdominal hair was shaved, and the skin was cleaned with chlorhexidine followed by povidone-iodine solution. A midline laparotomy incision of 2-3 cm was made to access the peritoneal cavity, and the intestine and colon were manipulated to expose the left renal artery. Vessels were separated from connective tissue and fat using blunt dissecting curved forceps and wet cotton swabs. The aorta was clamped above and below the renal artery bifurcation with microvascular clips. Next, both ends of the left renal artery were temporarily occluded using silk suture knot. A 1 mm incision was made on the occluded renal artery, a sterile device was inserted, and the incision was closed with tissue adhesive. The knot was opened first from the lateral, followed by the medial end. Clamps were released to allow kidney reperfusion, ensuring that the arterial occlusion did not exceed 15-20 min. The area was checked for active bleeding and observed for an additional 10-15 min. The abdominal wall was closed in two layers with 4-0 absorbable sutures using a simple continuous pattern for the muscle and a simple interrupted pattern for the skin. Topical ointment was applied to the suture site, and the animal was returned to its cage on a heating pad until fully recovered. The total surgery time ranged from 30 to 45 min. The internal diameter of the renal artery in rats was approximately 1.1 mm with a cross-sectional area of 0.95 mm^2^. For cohort 1, the diameter of the helical-shaped device was approximately 1 mm with a cross-sectional area of approximately 0.62 mm^2^ and a lumen diameter of 0.45 mm. For cohort 2, the dimensions of the linear-shaped devices were approximately 0.4 mm x 0.25 mm with a cross-sectional area of 0.1 mm^2^. Post-surgery, animals received meloxicam (2 mg/kg s.c.) every 24 h for 72 h. Blood collection for serum drug level estimation was performed via retro-orbital puncture on day 7 under anesthesia with isoflurane. The serum was separated by centrifugation (6000 rpm, 10 min, 4 °C). The animals were sacrificed using pentobarbital and phenytoin cocktail (Euthasol) (200 mg/kg) and the left and right kidneys were isolated, homogenized in ice-cold phosphate buffer (10% w/v), centrifuged (10000 rpm, 10 min, 4 °C), and the supernatants were collected for DEX quantification using LC-MS/MS as described previously. Liver, heart, lungs, and spleen were also collected from rats in the Kidney Cohort 2 with linear IADD devices. Implanted and control renal arteries were isolated and stored in 4% paraformaldehyde in phosphate-buffered saline (PBS) for further analyses.

#### 2.8.2 Focal delivery of DEX to the brain in rats

Fischer rats were divided into two groups: Brain-IADD (n=5) and Brain-Control (n=3). The Brain-IADD group had linear-shaped DEX-loaded IADD devices (fabricated by the two-step method) surgically implanted in the right carotid artery for focal drug delivery to the brain. The Brain-Control group received DEX (5 mg/mL) in drinking water.

The rats were prepared and anesthetized using the similar procedures as described above for the kidney cohorts. A pre-operative subcutaneous injection of 10 mL/kg normal saline was given before placing the animal in a supine position with its limbs fixed to the surface using low-tack adhesive, ensuring the upper extremities remained in a normal position to prevent lung compression. Once the depth of anesthesia was confirmed by monitoring reflexes such as withdrawal from a toe pinch, the hair over the incision site was shaved with a sterile razor blade, and the skin was cleaned with chlorhexidine followed by povidone-iodine solution. The site was then infused with bupivacaine. For common carotid artery implantation, an incision was made on the neck to expose the common carotid artery by dissecting the submandibular glands and sternohyoid muscles. The artery was occluded using vascular clamps. A 1-mm incision was made on the artery, a sterile device was inserted, and the incision was closed using tissue adhesive. The carotid artery was occluded for approximately 15 mins. The device self-adhered to the wall of the artery owing to its property of tissue adhesion. After removing the clamps and ensuring there was no internal bleeding along with no visual displacement of the device within the artery, the skin incision was closed using simple interrupted sutures. A topical ointment was applied to the suture site, and the animal was returned to its cage on a heating pad until fully recovered. The total surgery time, from incision to suture, was approximately 30-45 min. The internal diameter of the carotid artery in rat was approximately 1.1 mm with a cross-sectional area of 0.95 mm^2^. The dimensions of our linear devices were approximately 0.4 mm x 0.25 mm with a cross-sectional area of 0.1mm^2^. Post-surgery, the animals received meloxicam (2 mg/kg s.c.) every 12-24 h for 72 h. To determine serum drug level, blood collection was performed via retro-orbital puncture on day 7 under isoflurane anesthesia. The serum was separated by centrifugation (6000 rpm, 10 min, 4 °C). Animals were sacrificed using pentobarbital and phenytoin cocktail (Euthasol) (200 mg/kg) and the brain and liver were isolated, homogenized in ice-cold phosphate buffer (10% w/v), centrifuged (10000 rpm, 10 min, 4 °C), and the supernatants collected for DEX quantification using LC-MS/MS as described previously. The liver was included in this experiment as an indicator for off-target delivery to another organ. Implanted and control carotid arteries were isolated and stored in 4% paraformaldehyde in PBS for further analyses. To assess longer-term vascular and downstream brain tissue responses after carotid implantation, an additional small cohort of Fischer rats was implanted with linear one-step DEX-loaded IADD devices in the right carotid artery and examined for 55 days after implantation (n = 2). The surgical implantation procedure was performed as described above for the 7-day carotid implantation cohort. At the 55-day endpoint, animals were euthanized, and the implanted carotid artery region was isolated for histological assessment. The contralateral unimplanted carotid artery was collected and used as an internal control. Brains were rapidly collected and sectioned coronally using a brain matrix for 2,3,5-triphenyltetrazolium chloride (TTC) staining to assess macroscopic evidence of ischemic injury. Brain sections were incubated in 2% TTC solution at 37 °C for 30 min, rinsed, and imaged using a flatbed scanner (EPSON XP-7100) under consistent scanning settings. Arterial tissues were fixed and processed for Martius Scarlet Blue staining (MSB) to assess thrombus/fibrin deposition and local vascular morphology after prolonged implantation.

To quantify relative focal targeting, we determined the normalized organ drug levels (NODL), using the equation below:

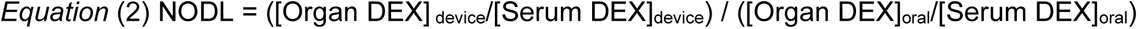

NODL was used as a relative index of focal targeting efficiency rather than as a dose-matched pharmacokinetic comparison. This metric is most applicable for comparing the extent of target-organ enrichment relative to systemic exposure within each route of administration. Because the IADD and oral groups differed in total administered dose, absorption kinetics, and route-dependent bioavailability, NODL does not directly estimate absolute exposure, total delivered dose, or therapeutic efficacy. For samples below the lower limit of detection (LLOD), the LLOD value was assigned conservatively for calculation purposes. This conservative assignment was used to avoid overestimating focal targeting when serum DEX was undetectable in IADD-treated animals; however, it may underestimate the true NODL when actual serum concentrations are below the LLOD.

#### 2.8.3 Histopathology of implanted arteries

Staining for histopathology was performed as described with slight modifications [40]. Briefly, arteries (renal and carotid) were fixed in 10% neutral buffered formalin, processed, and embedded in optimum cutting temperature (OCT) medium. Sections of 10-µm thick were stained in Mayer’s hematoxylin (3-5 min), rinsed, and “blued” in a mildly alkaline solution (10-15 s). Sections were counterstained with eosin Y (30 s) and cover-slipped with a resinous medium.

For the 55-day carotid implantation cohort, MSB staining was performed on arterial sections to evaluate thrombus and fibrin deposition around the implantation site. Staining was performed according to the manufacturer’s protocol (VitroView Martius Scarlet Blue Stain Kit, Vitro Vivo Biotech LLC). Sections were evaluated qualitatively for luminal patency, thrombus/fibrin deposition, vessel morphology, and local tissue response.

### 2.9 Statistical Analysis

All sets of data were expressed as Mean±SD and analyzed using GraphPad Prism v10.2.3 (GraphPad Prism software, San Diego, CA, USA). IC_50_ values were calculated with non-linear regression (curve fit) method as the drug concentration at the 50% cell viability. Comparison of in vivo drug release data was performed using one-way analysis of variance (ANOVA) followed by Tukey’s post-hoc comparisons. Normality was verified using the Shapiro-Wilk test, and the datasets met the assumptions for parametric analysis. The comparison of DEX levels between the brain and serum levels was performed using the paired t-test. The difference in the means for all the tests was considered statistically significant at *p*<0.05.

## 3. Results

The results of this work are to demonstrate proof-of-concept for our novel intra-arterial drug delivery (IADD) devices by evaluating the design and fabrication processes, *in vitro* release, cytocompatibility, released-drug bioactivity, and *in vivo* biodistribution in rat models.

### 3.1 Microstructure and Composition of Drug-loaded IADD Devices and Controls

We designed and fabricated IADD devices and systematically characterized their microstructure and composition to ensure reproducibility and consistency across batches (Fig. 1). We tested the viscosity of pre-PGS from 0°C to 120°C (Fig. 1A). The viscosity of pre-PGS used for IADD device fabrication was similar to previously reported viscosity profiles [24]. For this study, the PGS was synthesized with reduced time to have a lower viscosity, thus achieving a uniform PGS (or drug-loaded PGS) coating onto Mg substrates in the IADD devices. Model IADD devices with two different geometries were used, a helical shape (Fig. 1B) and a linear plank structure (Fig. 1C). For the helical design, the gaps between the Mg coils were completely filled with drug-loaded PGS coating, resulting in a solid tubular structure. For the linear design, the Mg wire was centrally positioned and surrounded with drug-loaded PGS coating. Further examination of the device cross-section in SEM (Fig. 1E-G) confirmed that the Mg microwire was fully embedded within the PGS matrix. SEM-EDS analysis was used to further assess the elemental composition of the PGS-Mg devices after drug loading (Fig 1H). The PGS control was composed primarily of carbon and oxygen, consistent with the polymer matrix. DEX-loaded PGS devices prepared by both two-step and one-step loading showed similar carbon- and oxygen-rich spectra, with low but detectable fluorine signal (0.14–0.15%), consistent with the presence of fluorinated DEX within the PGS matrix. In contrast, CIS-loaded PGS devices showed detectable chlorine (1.10%) and platinum (0.68%) signals, consistent with incorporation of cisplatin. Magnesium was detected in drug-loaded samples, reflecting contribution from the underlying Mg microwire core or exposed Mg regions within the analyzed device cross-section. Together, these SEM-EDS results support the presence of drug-associated elemental signatures in the PGS-Mg IADD devices, while confirming the expected carbon- and oxygen-rich composition of the PGS matrix. Notably, Pt and Cl signals were not detected in the EDS of DEX-IADD devices and no-drug control IADD, while F was not found in the EDS of CIS-IADD devices and no-drug control IADD, confirming the expected drug loading in the as-fabricated devices.

**Fig. 1.**
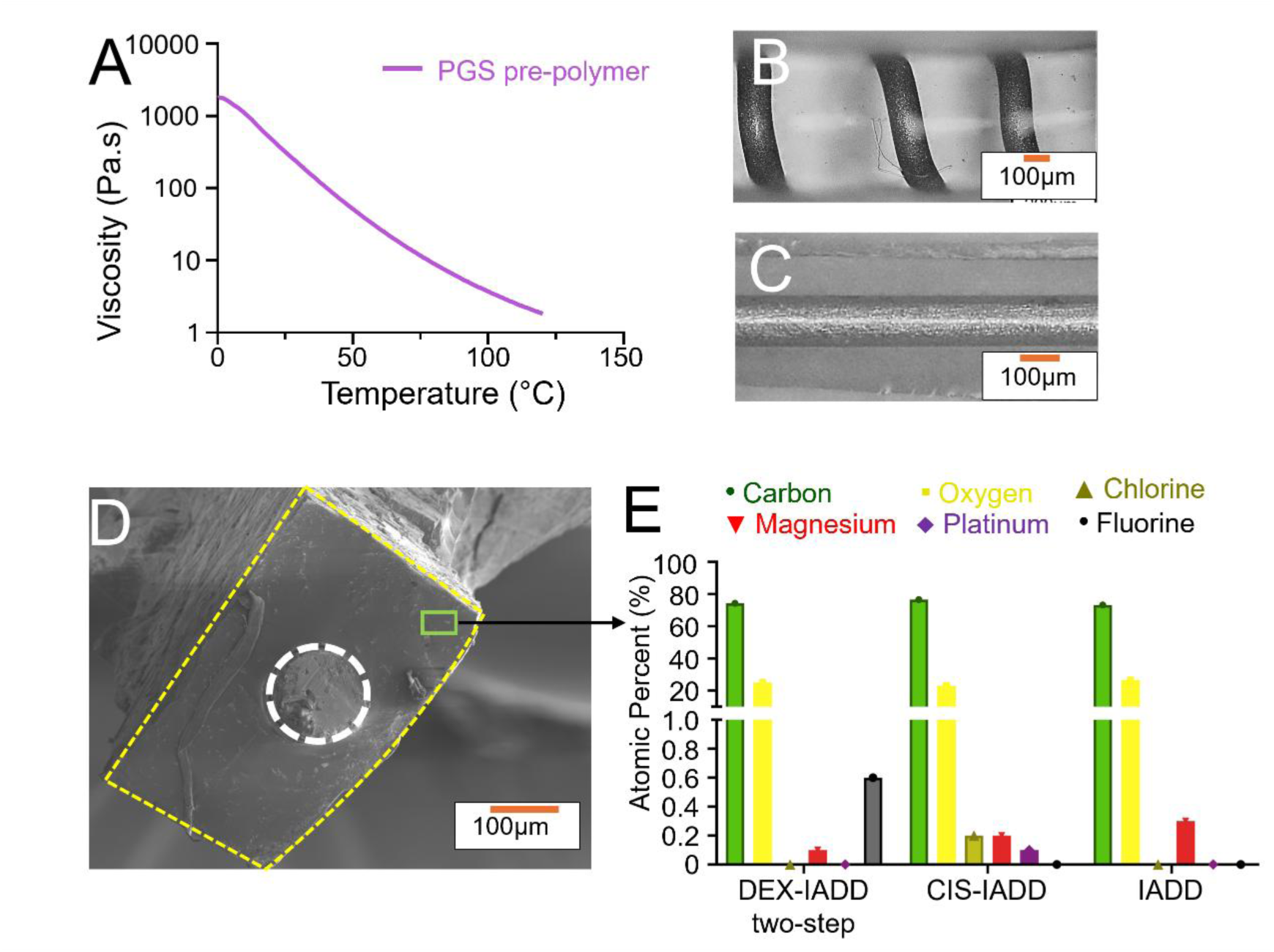
Structural and compositional characterization of drug-loaded IADD devices. A) Viscosity of PGS pre-polymer. B, C, D) Different geometries of IADD devices, (B) a helical shape (C, D) and linear structure. E, F ,G) SEM images of (E) DEX-IADD two-step, (F) DEX-IADD one-step, and (G) CIS-IADD. E) Elemental composition of drug-loaded IADD devices, which supports incorporation of the drug within the PGS matrix.

We further analyzed the drug-loaded and control devices using TGA and FTIR-ATR (Fig. 2). We tested the weight loss of drug-loaded IADD devices, IADD controls, DEX, and CIS, when the temperature increase caused the decomposition of PGS and drugs (Fig. 2A). PGS began decomposing at 394 °C and fully decomposed at 504 °C, leaving no solid residue. The DEX-only sample began decomposing at 285 °C and decomposed almost completely by 495 °C, leaving less than 2% solid residue, likely carbon black from incomplete combustion. The CIS-only sample began decomposing at 315 °C and continued to 364 °C; and, the weight loss then reached a plateau after decomposition, with approximately 65% mass remaining, because Pt in CIS has a high melting temperature of 270 °C. The DEX-loaded and the control IADD devices showed similar heat decomposition since the DEX amount is small relative to the PGS matrix. In the CIS-loaded IADD devices, 3% residue remained, consistent with Pt as a solid remnant after the CIS decomposition. FTIR-ATR spectra of drug-loaded IADD devices were compared with non-drug-loaded IADD controls and drug controls (Fig. 2B). The FTIR spectra of drug-loaded IADD devices were highly similar to IADD controls and revealed evidence of drug incorporation without chemical degradation. The PGS in the IADD device exhibits its characteristic ester carbonyl stretch (∼1730-1740 cm^-1^), along with broad O-H stretching and C-O bands in the 1000-1300 cm^-1^ region, similar to the FTIR-ATR spectra of PGS reported in the literature [26]. Upon loading DEX or CIS, the composite spectra of DEX-IADD and CIS-IADD retained the core polymer peaks, with attenuated and slightly shifted drug-related features. The FTIR-ATR spectra of DEX-loaded IADD devices and control devices are similar, and no significant difference was observed, considering the low amount of drug relative to the PGS matrix and overlap of peaks between IADD controls and DEX. In the CIS-IADD spectrum, weak N-H stretching (∼3200-3400 cm⁻¹) and amide bending (∼1600-1650 cm⁻¹) bands remained discernible, although diminished relative to pure CIS. The absence of new peaks and the preservation of polymer backbone bands confirmed the drug encapsulation in PGS without covalent modification or decomposition.

**Fig. 2.**
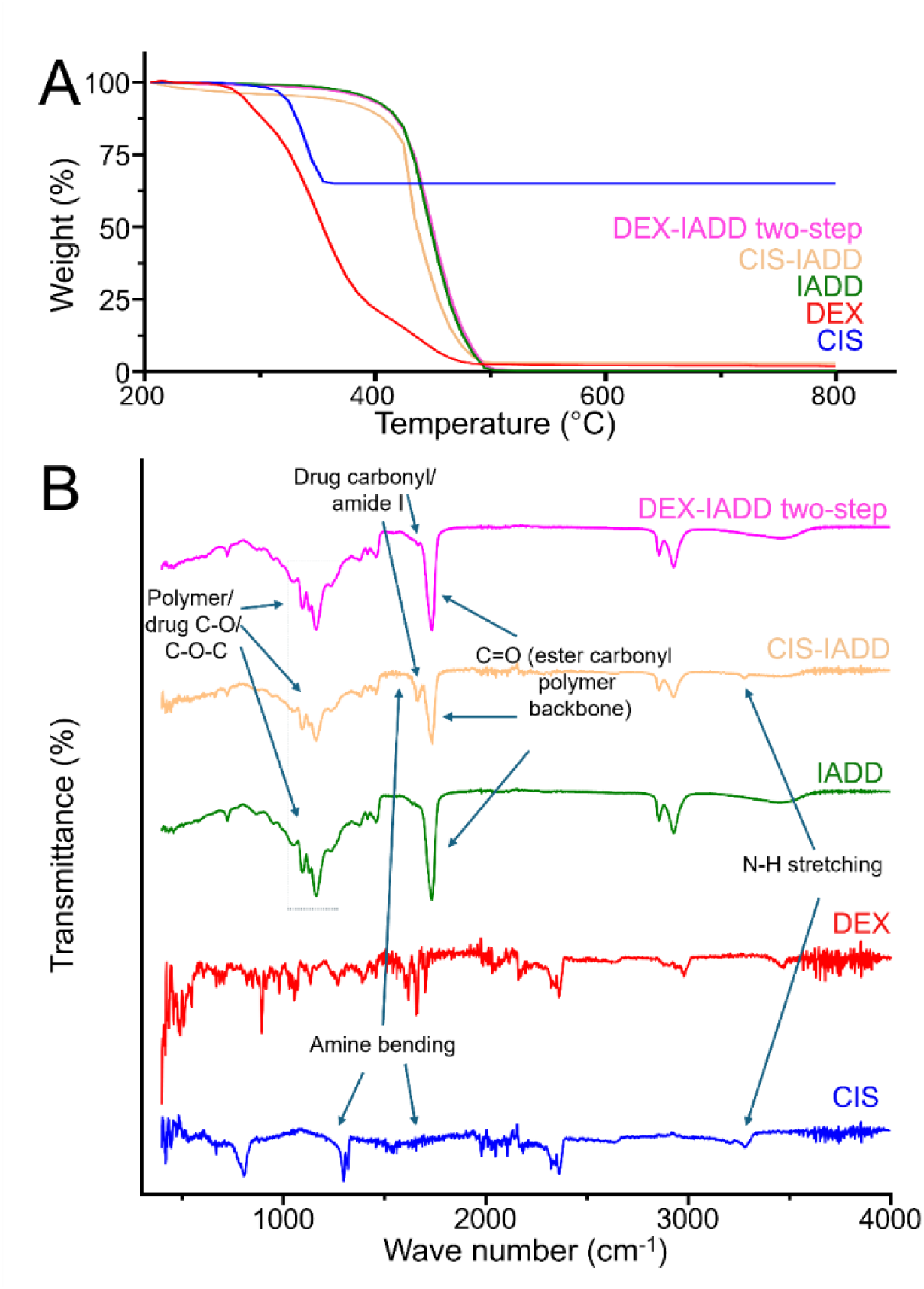
Thermal and spectroscopic characterization of drug-loaded IADD devices. A) TGA curves of DEX- and CIS-loaded IADD devices in comparison with non-drug-loaded IADD device and standard model drugs of DEX and CIS. B) FTIR-ATR spectra of DEX- and CIS-loaded IADD devices in comparison with non-drug-loaded IADD device and standard model drugs of DEX and CIS.

### 3.2 Drug Release Profiles and Degradation of DEX-Loaded and CIS-Loaded IADD Devices In Vitro

Building on the successful fabrication of the IADD devices, we next quantified their drug release profiles in physiological solutions and device degradation *in vitro* over 30 days (Fig. 3). The amounts of absolute and cumulative DEX and CIS released from DEX-loaded and CIS-loaded IADD devices over 30 days, with daily medium replacement, are presented (Fig 3A and 3B). The DEX-loaded IADD devices released cumulative 373.11 ± 1.41 µg of DEX over the 30 days, with a burst release on day 1 (274.65 ± 91.90 µg), equivalent to 73% of total release. By day 7, DEX release stabilized between 0.39 - 4.12 µg/day with an average release of 1.78 ± 1.12 µg/day. CIS-loaded IADD devices did not demonstrate a burst release phase and released a cumulative 64.73 ± 0.06 µg of CIS over 30 days. Similar to DEX, CIS release stabilized by day 7 between 0.97 - 7.07 µg/day with minor fluctuations, releasing an average of 1.72 ± 1.2 µg/day. Because both DEX- and CIS-loaded devices in this experiment used the same linear geometry and two-step loading method, the distinct release profiles likely reflect differences in drug-polymer interactions, drug diffusivity, and effects of the loading-solvent.

**Fig. 3.**
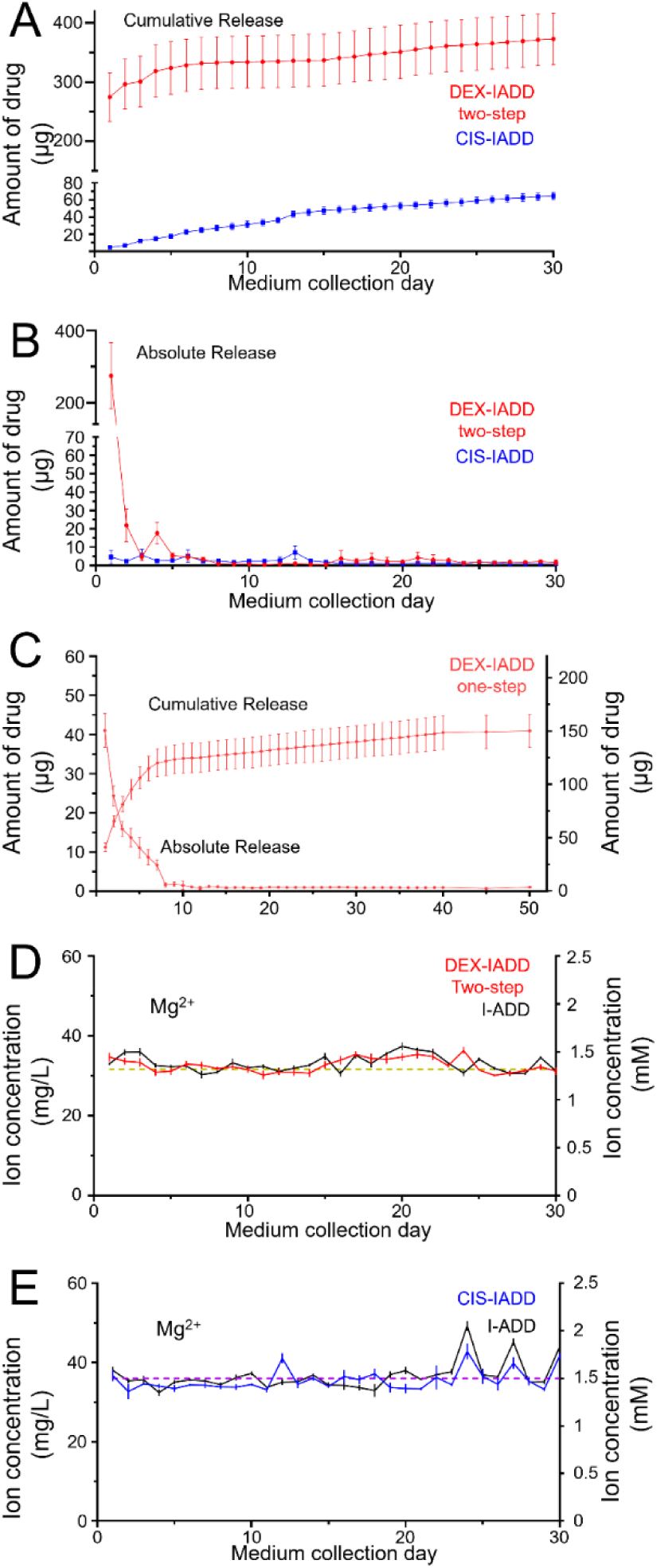
The drug release and magnesium ion release profiles in physiologically relevant solutions *in vitro*. A) Cumulative amount of DEX and CIS released from two-step DEX-loaded and CIS-loaded IADD devices per day for 30 days. B) Absolute amount of DEX and CIS released from two-step DEX-loaded and CIS-loaded IADD devices per day for 30 days. C) Absolute (left Y-axis) and cumulative (right Y-axis) amount of DEX released from one-step DEX-loaded IADD devices D, E) Magnesium ion concentrations in rSBF containing the DEX-loaded devices (D) and in aCSF containing the CIS-loaded devices (E), as well as the respective non-drug-loaded IADD control devices. Dashed lines reflect the baseline magnesium ion concentrations in the immersion media of rSBF or aCSF, that is, 1.3 mM (31.6 mg/L) in (D) and 1.5 mM (36 mg/L) in (E).

To further evaluate the one-step loading method, DEX release from one-step DEX-loaded IADD devices was monitored for 60 days in vitro. Quantifiable DEX release was observed through day 50, and undetectable post day 50, indicating measurable drug release for at least 50 days under these conditions (Fig 3C). The cumulative DEX recovered through day 50 was 150.06 ± 34.97 µg, corresponding to 94.18 ± 18.07% of the DEX introduced during fabrication. The day 1 release was 41.03 ± 9.70 µg, corresponding to 27.92 ± 5.93% of the cumulative DEX recovered through day 50. Compared with the two-step IADD device, these findings suggest that the one-step loading method reduced the relative burst release while maintaining prolonged DEX elution. To compare release behavior across loading methods, we summarized 30-day release metrics for two-step DEX-IADD, one-step DEX-IADD, and two-step CIS-IADD devices (Supplementary Table 1).

To evaluate the device degradation over time, Mg²⁺ ion concentrations in the collected media were quantified (Fig. 3D, E), as degradation of the Mg wire substrate releases Mg^2+^ ions. The measured Mg²⁺ ion concentrations demonstrated stable ion release throughout the 30-day period, without evidence of burst release that would indicate rapid or uncontrolled degradation. For DEX-loaded and non-drug-loaded IADD devices incubated in rSBF, Mg²⁺ ion concentrations remained near the baseline of rSBF controls at 31.6 mg/L (1.3 mM) (Fig. 3D). Similarly, for CIS-loaded and non-drug-loaded IADD devices incubated in aCSF, Mg²⁺ ion concentrations remained near aCSF controls of 36 mg/L (1.5 mM) (Fig 3E). These results suggest that the PGS coating substantially delayed Mg substrate degradation during most of the 30-day in vitro release period. Mg²⁺ concentrations did show modest increases in the later phases of incubation. These late-stage increases indicate increased degradation and permeability of PGS layer for water diffusion and ion release. However, the absence of an abrupt Mg²⁺ spike suggests that rapid or uncontrolled Mg degradation did not occur.

### 3.3 Cytocompatibility of Drug-loaded IADD Devices with HUVECs

Given the intended intra-arterial application of the IADD devices, we next evaluated their cytocompatibility with vascular endothelial cells (HUVEC) (Fig. 4). We determined the inhibitory concentration (IC_50_) profiles of the two drugs (DEX and CIS) and their respective solvents of ethanol and DMF (Fig. 4A, B). As expected, the drugs and solvents exhibited reduced cell viability at high concentrations. Importantly, these concentrations were substantially higher than those expected to be released from our IADD devices under physiological conditions. We assessed the cytocompatibility of the drug-loaded IADD and control devices with HUVECs using the exposure culture method (Fig. 4C). No significant differences in viability were observed across the groups of interest, including two-step DEX-loaded IADD (100.5 ± 1.57%), two-step CIS-loaded IADD (94.94 ± 0.37%), ethanol-IADD control (98.38 ± 2.11%), DMF-IADD control (101.0 ± 1.33%), and non-treated cell only control (100.0 ± 1.14%) (Fig. 4C). Collectively, these *in vitro* results suggested that the IADD devices are cytocompatible with endothelial cells for potential intra-arterial implantation *in vivo*, while also confirming effective removal of residual solvents from the IADD devices during fabrication.

**Fig. 4.**
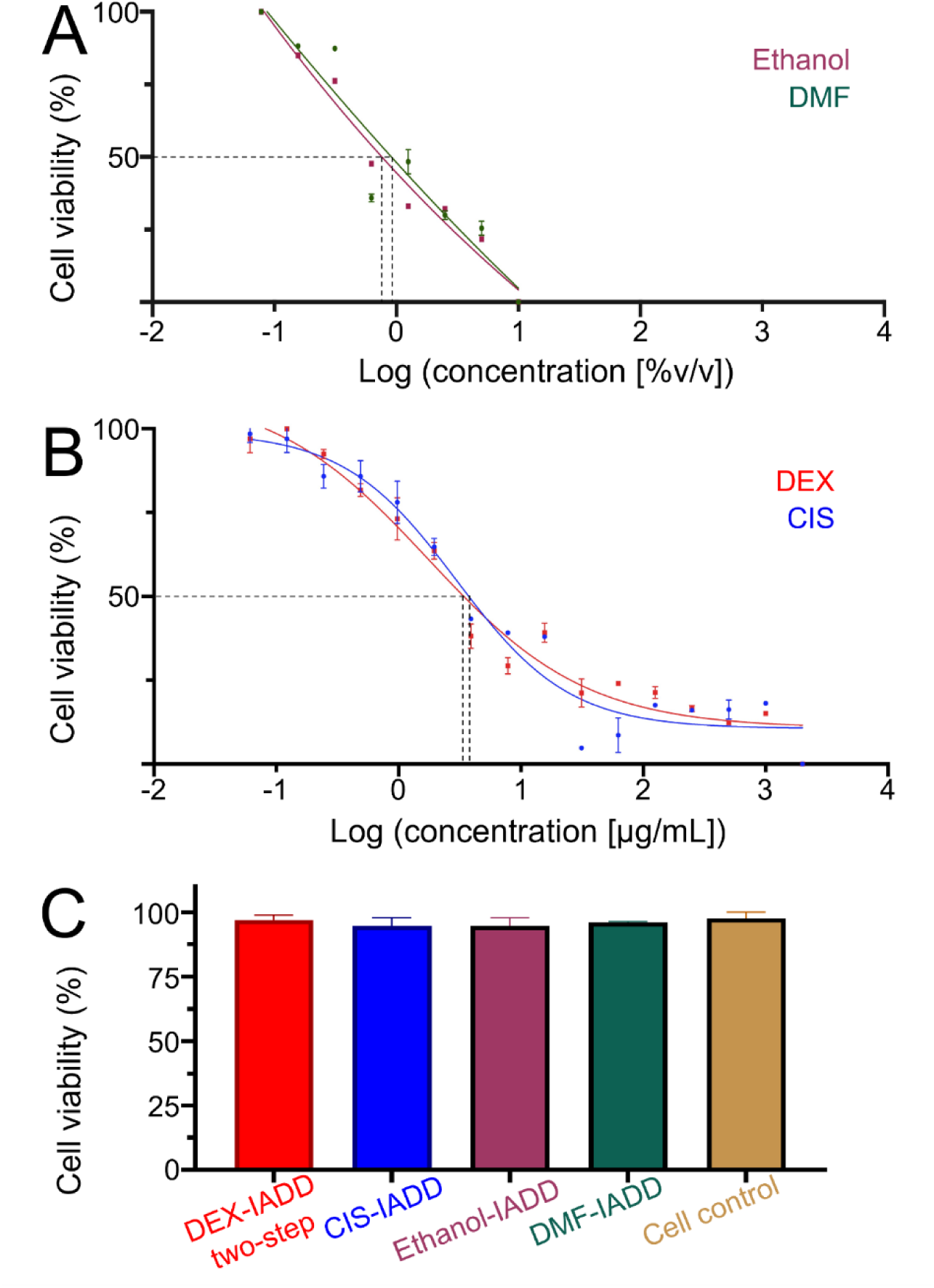
Cytocompatibility of drug-loaded IADD devices with HUVEC cells *in vitro* along with drug and solvent controls. A, B) IC_50_ curves of (A) DEX and CIS drugs and (B) ethanol and DMF solvents. C) Percentages of HUVEC viability when exposed to the two-step DEX-loaded, two-step CIS-loaded, ethanol-soaked, and DMF-soaked IADD devices in comparison with the cells only controls.

### 3.4 CIS-loaded IADD Devices Reduced Viability of Glioma Cells *In Vitro*

To evaluate the pharmacological activity of the released drug, we examined the viability of F98 glioma cells exposed to the media collected daily from the two-step CIS-loaded IADD devices during the 30-day *in vitro* drug release. The inhibitory concentration (IC_50_) for CIS was determined as 2.21 µg/mL (Fig. 5A), confirming effective cytotoxic activity against F98 cells. When the cells were treated with the daily release media from two-step CIS-loaded IADD device, an average reduction in cell viability of 38.21 ± 10.56% was observed (Fig. 5B). These findings validated that the CIS released from the IADD devices remained pharmacologically active during prolonged release. Interestingly, F98 cells cultured with the daily release media from non-drug-loaded IADD controls exhibited higher cell densities and elevated viability compared to non-treated cell only controls (Fig. 5B). We speculate that the enhanced viability may be because the degrading PGS would release glycerol and the cells could use glycerol as an energy source [41, 42].

**Fig. 5.**
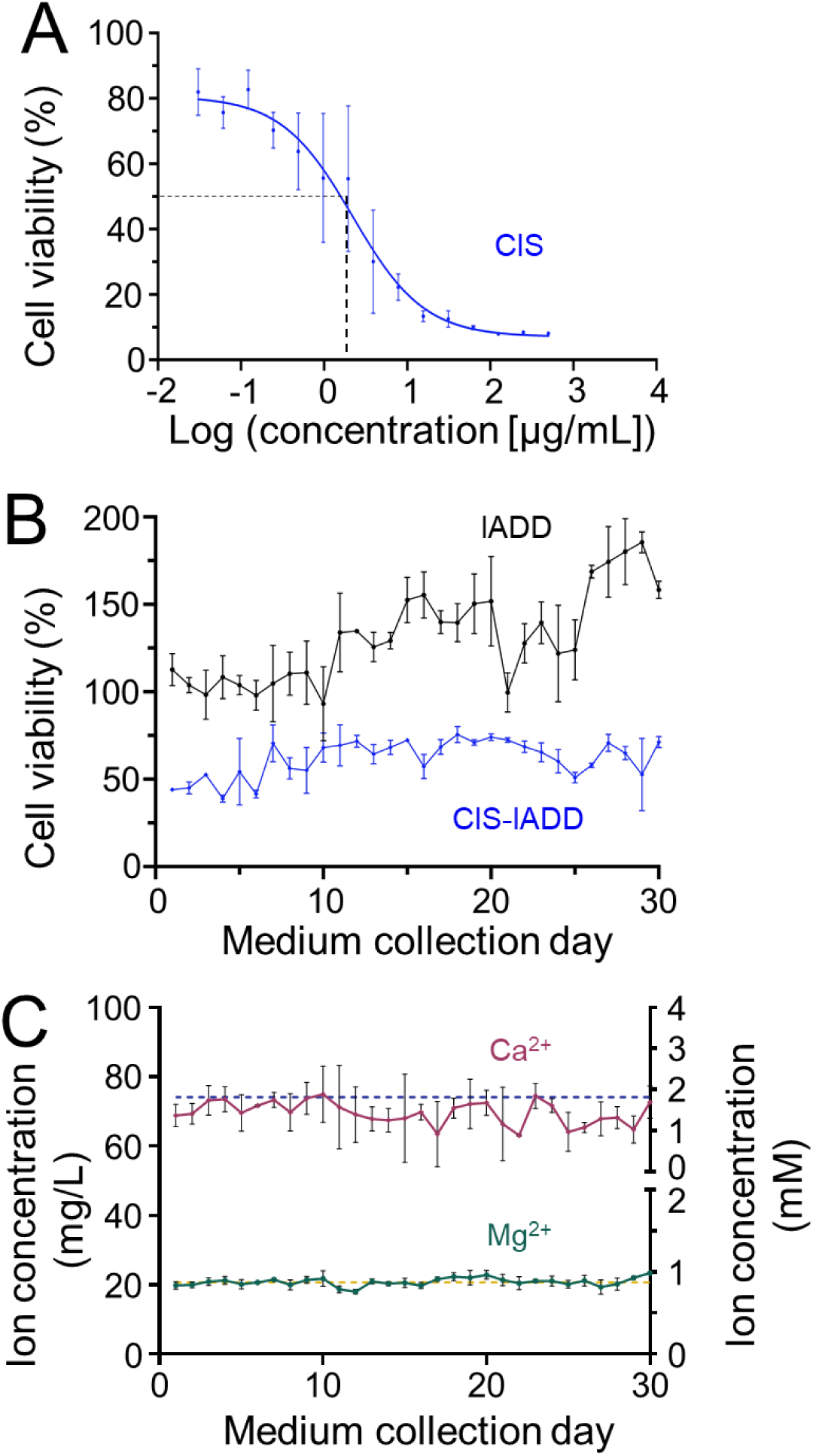
In vitro drug activity of CIS released from the two-step CIS-loaded IADD devices against F98 glioma cells. A) IC_50_ curve of CIS. B) Percentage of glioma cell viability when exposed to the release media collected from two-step CIS-loaded IADD device (blue) versus non-drug-loaded control device (black). The assay measures F98 viability after exposure to each day’s independently collected release medium, and ‘medium collection day’ refers to the day that the release media were collected. C) Mg^2+^ and Ca^2+^ ion concentrations in the culture media of the F98 cells doped with two-step CIS-loaded IADD release media (solid lines). Dashed lines reflect ion concentrations from culture media controls: 0.8 mM (19.4 mg/L) of Mg^2+^ (bottom right y-axis) and 1.8 mM (72.1 mg/L) of Ca^2+^ (top right y-axis).

To further assess the metabolic impact of the released drug, Mg²⁺ and Ca²⁺ ion concentrations were measured in the culture media of the F98 glioma cells exposed to two-step CIS-loaded IADD release media (Fig. 5C). These ions served as indirect indicators of cellular metabolic stability. Mg²⁺ and Ca²⁺ concentrations remained stable throughout the 30-day sampling period, with average levels comparable to those observed in the media control group. These findings suggest that the release media from two-step CIS-loaded IADD devices did not significantly alter Mg²⁺ or Ca²⁺ homeostasis in F98 cells, indicating preserved cellular metabolic balance during exposure.

### 3.5 DEX-loaded IADD Devices Demonstrated Focal Drug Delivery in Targeted Kidney and Brain *In Vivo*

The *in vivo* studies were designed to evaluate whether DEX-loaded IADD devices could increase target-organ drug levels relative to systemic exposure in two rat implantation models. Specifically, the DEX-loaded IADD devices with two different geometries and/or loading methods were implanted in the left renal artery of rats upstream of the left kidney, or in the right carotid artery of rats upstream of the brain. The devices with helical design (fabricated by the two-step method) were implanted into rat Kidney Cohort 1 (Fig. 6), the devices with linear design (fabricated by the one-step method) were implanted into rat Kidney Cohort 2 (Fig. 7), and the devices with linear design (fabricated by the two-step method) were implanted into rat brain cohort (Fig. 8). Because device geometry, loading method, drug payload, and implantation site were not independently varied, differences among these cohorts should be interpreted as prototype-specific outcomes rather than as isolated effects of any single design variable. All the animals survived during the 7-day period post implantation with the IADD devices. For comparison, oral control rats received DEX through their drinking water (1 mg/L or 5 mg/L). These concentrations were selected to produce pharmacologically relevant systemic DEX exposure in rats and are sufficient to induce adrenal suppression in short-term studies [43–45]. Across cohorts, DEX concentrations were higher in the intended downstream target organs than in serum or off-target compartments, supporting focal target-organ enrichment by IADD-mediated delivery. Histological analyses of explanted arteries provided preliminary evidence of local vascular tolerability over the examined implantation period.

**Fig. 6.**
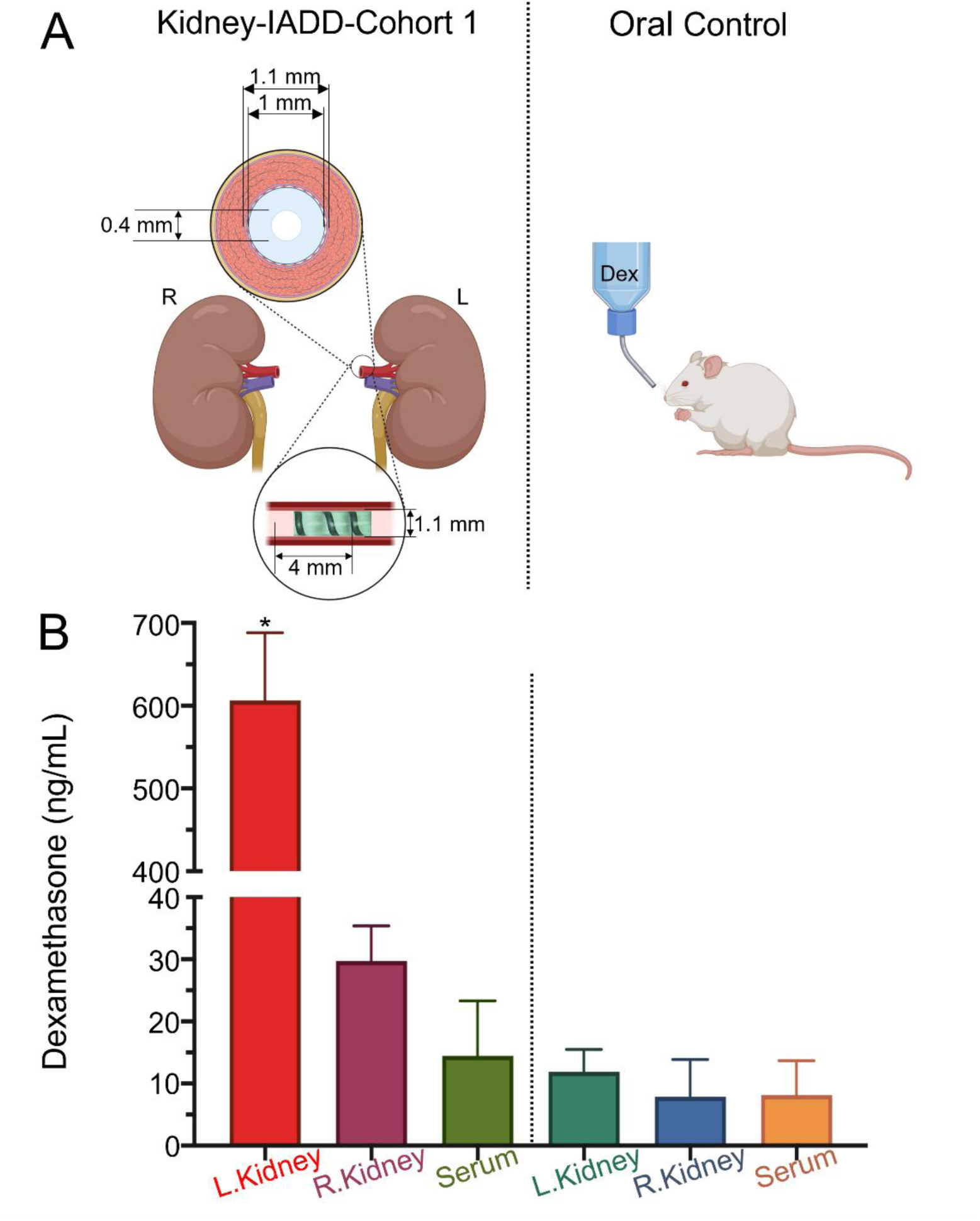
DEX-loaded IADD devices enhanced focal drug delivery in targeted kidney *in vivo* in the rat Kidney Cohort 1. A) Schematic representation of the *in vivo* drug administration via helical-shaped IADD device (left) or orally (right). B) DEX concentrations in the kidneys and the serum, after IADD device implantation in the left renal artery for 7 days (left) or after 7 days of oral DEX administration (right). \**p* < 0.05 compared to all other compartments in the IADD-treated animals.

**Fig. 7.**
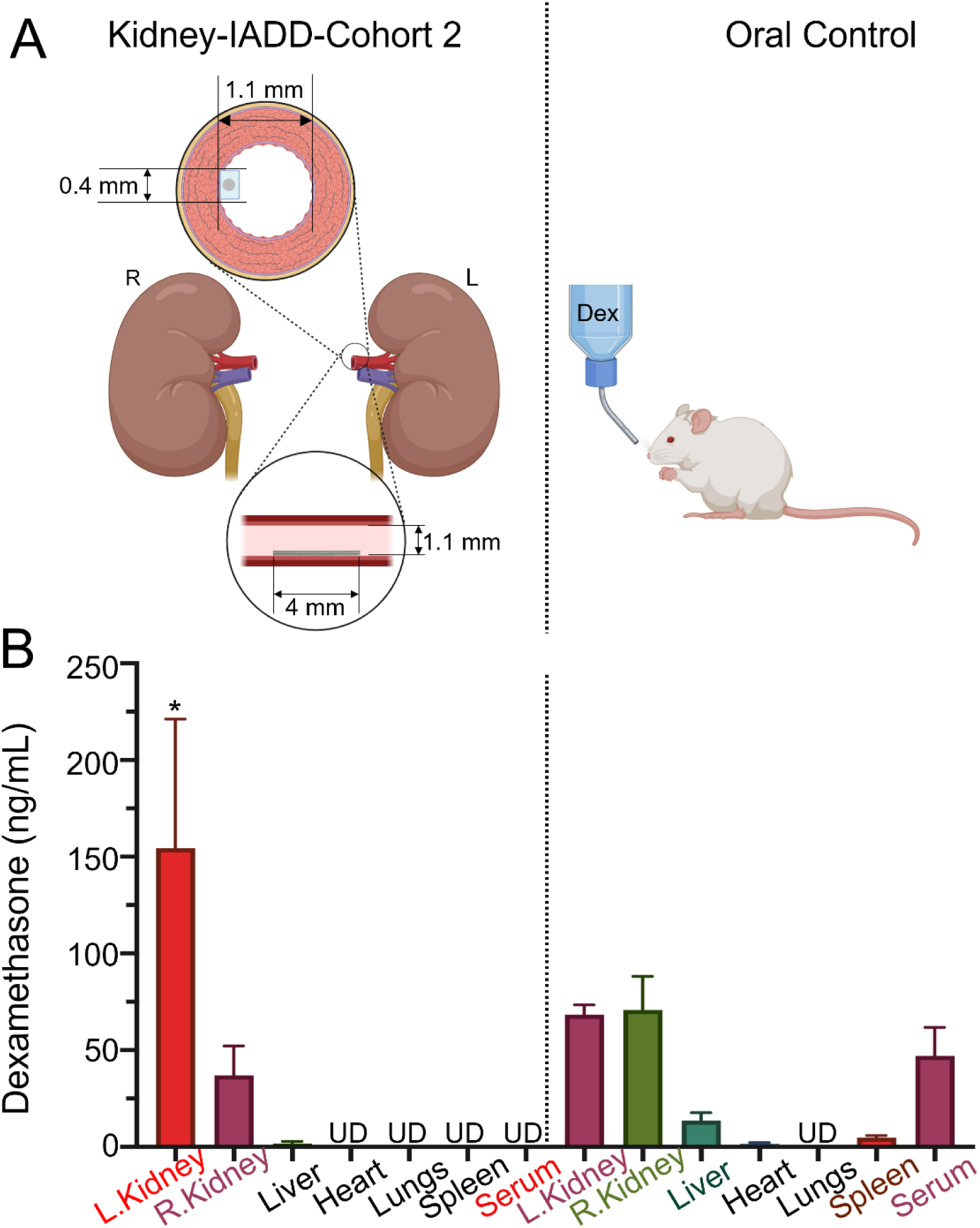
DEX-loaded IADD devices enhanced focal drug delivery in targeted kidney *in vivo* in the rat Kidney Cohort 2. A) Schematic representation of the *in vivo* drug administration via the linear-shaped IADD device (left) or orally (right). B) DEX concentrations in the left kidney (target), right kidney (contralateral), serum, and off-target organs (liver, spleen, heart, lungs) after IADD device implantation in the left renal artery for 7 days (left) or after 7 days of oral DEX administration (right). “UD” indicates values below the LC-MS/MS LLOD. \**p* < 0.05 compared to all other compartments in the IADD-treated animals.

**Fig. 8.**
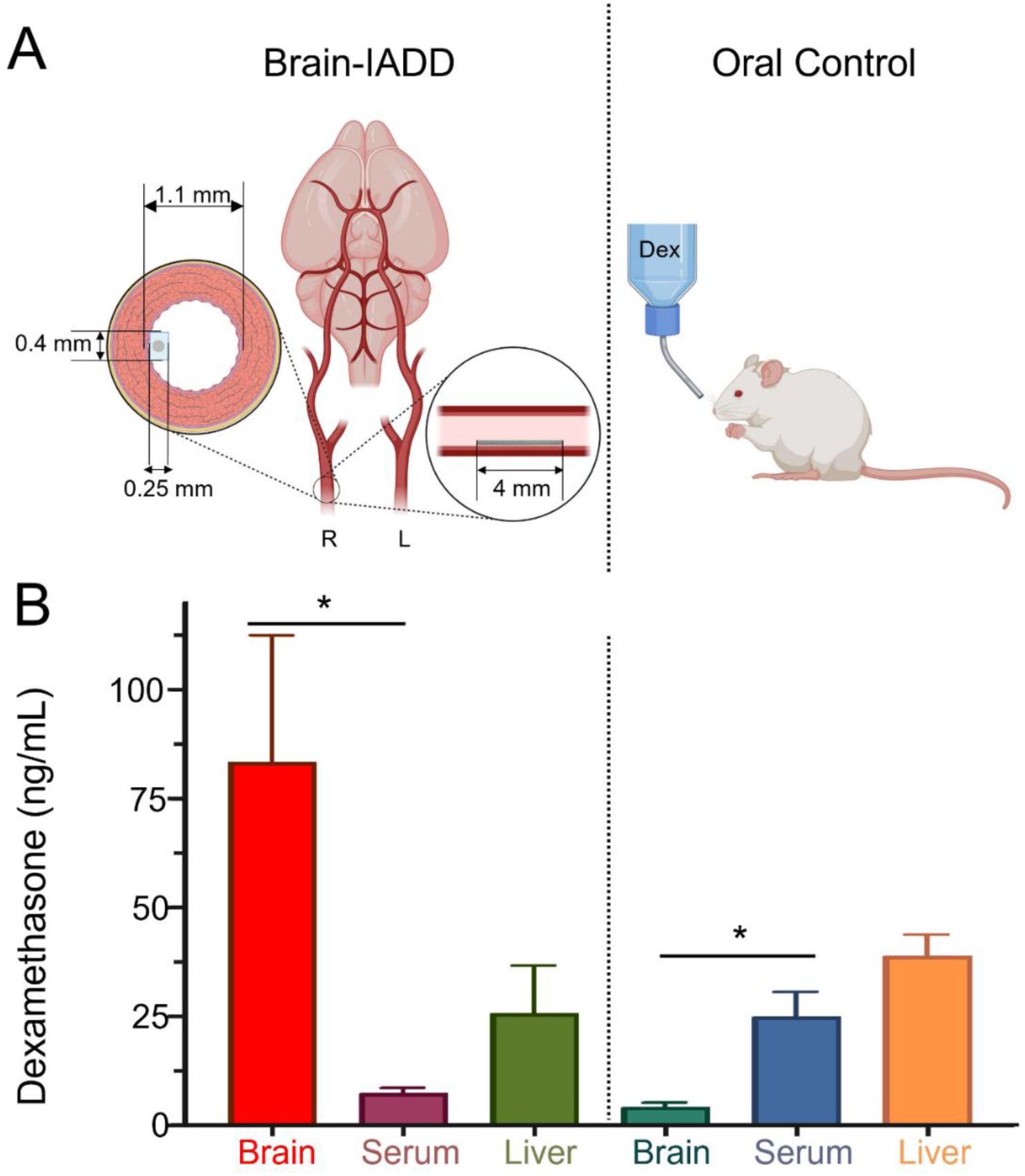
DEX-loaded IADD devices enhanced focal drug delivery in targeted brain *in vivo* in the rat Brain Cohort. A) Schematic representation of the *in vivo* drug administration via the linear-shaped IADD device (left) or orally (right). B) DEX concentrations in the brain, serum, and the liver, after IADD implantation in the right carotid artery for 7 days (left) or after 7 days of oral DEX administration (right). * *p*<0.05.

#### 3.5.1 DEX-loaded IADD devices increased DEX levels in targeted kidney *in vivo*

In the Kidney Cohort 1, the two-step DEX-loaded IADD devices (Fig. 6A) contained a larger volume of drug-loaded PGS per device because of the helical-shaped design and two-step methods. In rats implanted with the devices in the left renal artery, DEX concentrations in the left kidney (606.40 ± 81.74 ng/mL) were markedly higher than in the serum (14.40 ± 8.88 ng/mL) and the right kidney (29.75 ± 5.62 ng/mL) (Fig. 6B). This corresponded to a 42-fold higher DEX concentration in the targeted left kidney compared with serum. In contrast, oral DEX administration (1 mg/mL in drinking water) resulted in comparable concentrations across all compartments, including the left kidney (11.88 ± 3.60 ng/mL), right kidney (7.88 ± 5.99 ng/mL), and serum (8.08 ± 5.54 ng/mL), as expected for systemic drug distribution of oral administration (Fig. 6B). The calculated normalized organ drug level (NODL) was 29, reflecting a 29-fold enhancement in kidney-to-serum drug concentration using the IADD devices versus oral dosing. Given the very high drug levels observed in the left kidney, we hypothesized that partial saturation of renal uptake might have contributed to the off-target exposure in the right kidney. To address this, in the Kidney Cohort 2, smaller, linear-shaped IADD devices fabricated by the one-step method were implanted in the left renal artery.

In the Kidney Cohort 2, the smaller linear-shaped one-step DEX-loaded IADD devices reduced lumen occlusion (Fig. 7A). In this cohort, DEX concentrations in the left kidney reached 154.4 ± 38.54 ng/mL, approximately 2.25-fold higher than levels in the left kidney following oral administration (68.39 ± 2.92 ng/mL) (Fig. 7B). Within the IADD-treated animals, serum DEX levels were below the lower limit of detection (LLOD = 0.97 ng/mL), indicating minimal systemic exposure. Additionally, DEX concentrations in the left kidney were significantly greater than those in the right kidney (36.95 ± 8.77 ng/mL) and liver (1.59 ± 0.61 ng/mL), while no detectable DEX was found in other off-target organs including the heart, lungs, and spleen (Fig. 7B). In contrast, the animals with oral DEX administration showed uniform DEX distributions across the left kidney (68.39 ± 2.92 ng/mL), right kidney (70.74 ± 10.07 ng/mL), and serum (47.02 ± 8.49 ng/mL), suggesting systemic drug exposure (Fig. 7B). For statistical analysis, undetectable concentrations were conservatively treated as LLOD. The calculated NODL for the Kidney Cohort 2 was 109, representing a 109-fold enhancement in focal drug targeting using the IADD device with linear design versus oral delivery. This improvement, substantially greater than the 29-fold enhancement achieved using the IADD device with helical design, indicates reduced drug loading and saturation in renal uptake for better local dosing efficiency. However, because Kidney Cohorts 1 and 2 differed in device geometry, loading method, and drug payload, the higher NODL in Cohort 2 cannot be attributed to a single design variable. Rather, these findings suggest that IADD design and loading parameters can influence focal biodistribution and should be systematically optimized per treatment condition.

#### 3.5.2 DEX-loaded IADD devices increased DEX levels in targeted brain *in vivo*

Two-step DEX-loaded IADD devices with the linear geometry significantly increased drug levels in the brain when implanted in the common carotid artery of rats (Fig. 8). In the device-treated animals, the DEX concentrations in the brain (83.46 ± 29.04 ng/mL) were significantly higher than in the serum (7.34 ± 1.24 ng/mL) (Fig. 8B). This 11-fold increase in brain versus serum concentration is particularly noteworthy given the restrictive nature of the blood-brain barrier. This is illustrated in the rats treated with oral administration, in which DEX levels in the brain (4.14 ± 1.05 ng/mL) were significantly lower than in the serum (24.95 ± 5.68 ng/mL). Furthermore, DEX concentrations in the brain from the IADD-treated animals were approximately 20-fold higher than those treated with oral dosing. To assess off-target distribution, the DEX concentrations were quantified in the liver (Fig. 8B). The DEX levels in the liver of IADD treated rats (25.74 ± 10.93 ng/mL) were comparable to that in rats with oral administration (38.93 ± 4.86 ng/mL), despite the 20-fold increase in brain DEX concentration. Based on these data, a normalized organ drug level (NODL) of 68 was calculated, indicating a 68-fold improvement in brain-specific targeting using IADD-mediated focal drug delivery versus oral administration. Together with the renal artery implantation studies, these data support the ability of DEX-loaded IADD devices to increase drug levels in downstream target organs relative to systemic exposure.

#### 3.5.3 Histopathology of explanted arteries and characterization of IADD Devices Post Implantation

After 7 days of implantation, histopathological changes in the explanted arteries were compared with non-implanted control arteries to assess the local vascular response to the IADD devices (Fig. 9). Hematoxylin and eosin (H&E) staining of representative left renal artery and right carotid artery sections showed patent lumens with preservation of the intima, media, and adventitia at the implantation sites. (Fig. 9B,D). Limited peri-arterial inflammatory infiltrates and focal adventitial remodeling were observed near the arteriotomy and adhesive interfaces, consistent with mild early post-surgical changes. No occlusive thrombus formation or transmural necrosis was identified in any of the examined fields, supporting the local vascular tolerability at the 7-day endpoint.

**Fig. 9.**
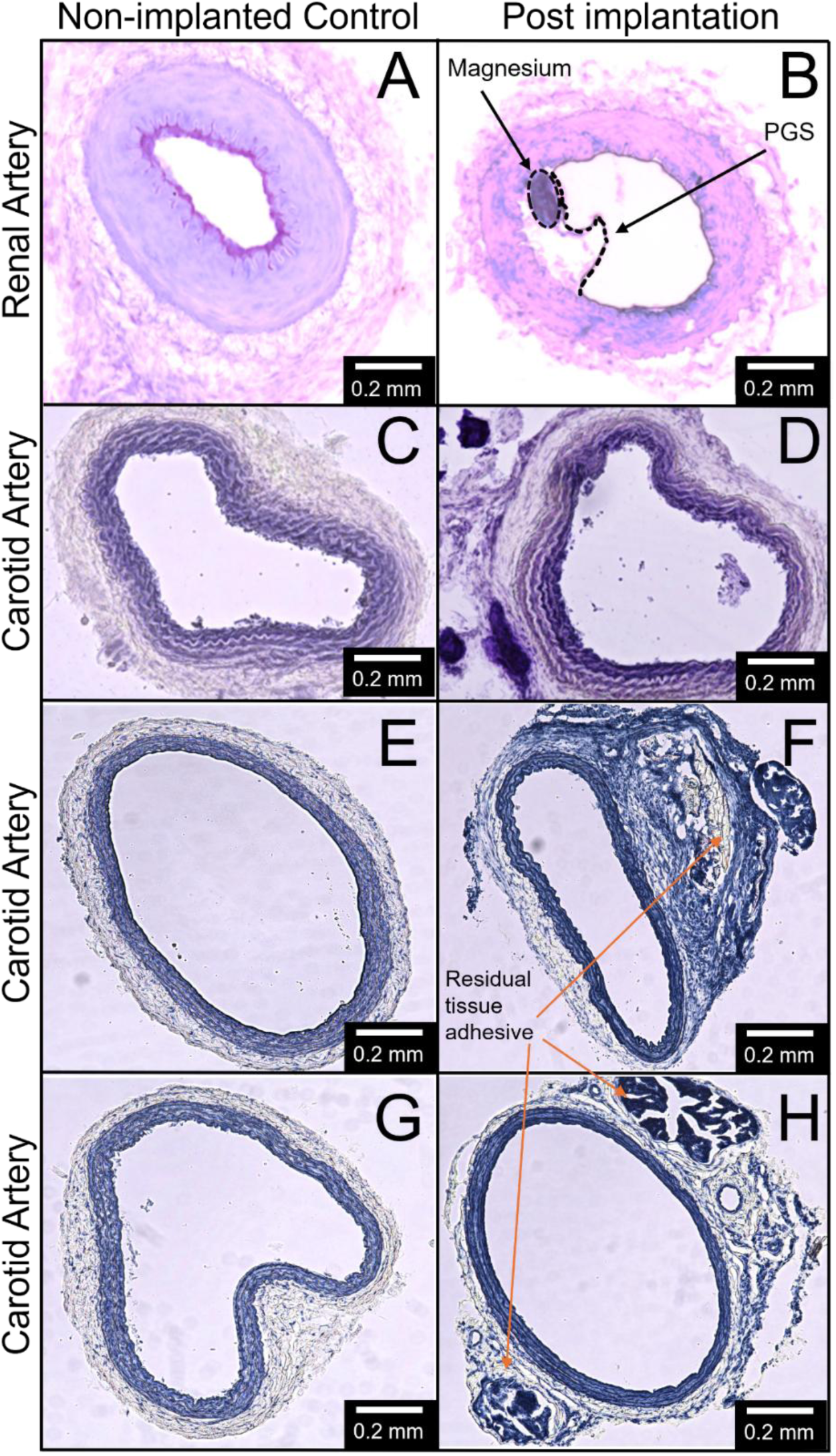
Histopathological analyses of explanted arteries. A-D) Representative H&E images of renal (A, non-implanted; B, explanted) and carotid (C, non-implanted; D, explanted) arteries. A fragment of the IADD device is observable in (B), showing the magnesium core (dashed circle) and surrounding PGS. The shape of the PGS protrusion is an artifact of sectioning. For the carotid artery in (D), the device dislodged during staining. E-H) Representative MSB-stained sections of carotid arteries collected at the 55-day endpoint, including implanted and contralateral unimplanted control arteries. MSB staining showed preserved arterial morphology and patent luminal architecture in the implanted carotid artery sections, with an appearance comparable to the contralateral unimplanted control arteries. No occlusive thrombus or major adverse vascular remodeling was observed in the examined sections. Residual tissue adhesive was noted outside the arterial wall and did not appear to compromise the arterial lumen.

To further extend the implantation interval, a separate small cohort of rats was examined 55 days after carotid artery implantation with one-step DEX-loaded IADD devices (n = 2). At this longer time point, the implanted device was no longer grossly evident in the examined arterial region. Arterial histology of implanted arteries showed preserved vessel morphology and patent luminal architecture (Fig. 9F,H), with an appearance comparable to the contralateral unimplanted control artery (Fig. 9E, G) and without evidence of occlusive thrombus or major adverse vascular remodeling in the examined sections after MSB staining. Residual tissue adhesive was noted outside the arterial wall in implanted arteries. To assess downstream brain tissue after prolonged carotid implantation, TTC staining was performed on brain sections from the same animals. No gross TTC-negative infarct regions were observed in the examined brains, suggesting absence of large territorial ischemic injury at the 55-day endpoint (Supplementary Fig. S2). These findings provide preliminary supportive evidence of longer-term local vascular tolerability and downstream brain tissue preservation after DEX-IADD implantation, although larger cohorts and quantitative vascular and neural endpoints will be required for definitive chronic safety assessment.

7-day explanted IADD devices were examined and compared to non-implanted control devices to assess their structural integrity following *in vivo* exposure (Fig. 10). In explanted devices, SEM-EDS analyses revealed partial surface degradation and interfacial modifications at the Mg-PGS boundary (Fig. 10D-F). The polymer layer remained largely intact but exhibited localized thinning and minor surface cracking, suggesting gradual surface erosion rather than bulk degradation. Elemental mapping confirmed that Mg remained in the device core (Fig. 10I), while an increased oxygen signal near the interface indicated the formation of magnesium oxide or hydroxide during physiological exposure (Fig. 10E, F, H). The polymer-rich regions contained predominantly carbon (C, 76.7-76.9 at.%) and oxygen (O, 22.4-22.5 at.%), which were similar in explanted and non-implanted devices (Fig. 10G, H). Fluorine peaks (F, ∼0.4 At%) were present in the polymer-rich regions of the explanted devices (Fig. 10J), indicating retention of DEX. Overall, the post-explantation profiles indicate limited surface oxidation and degradation of the Mg substrate and preservation of the polymer coatings, confirming the structural stability of the IADD devices after 7 days *in vivo*.

**Fig. 10.**
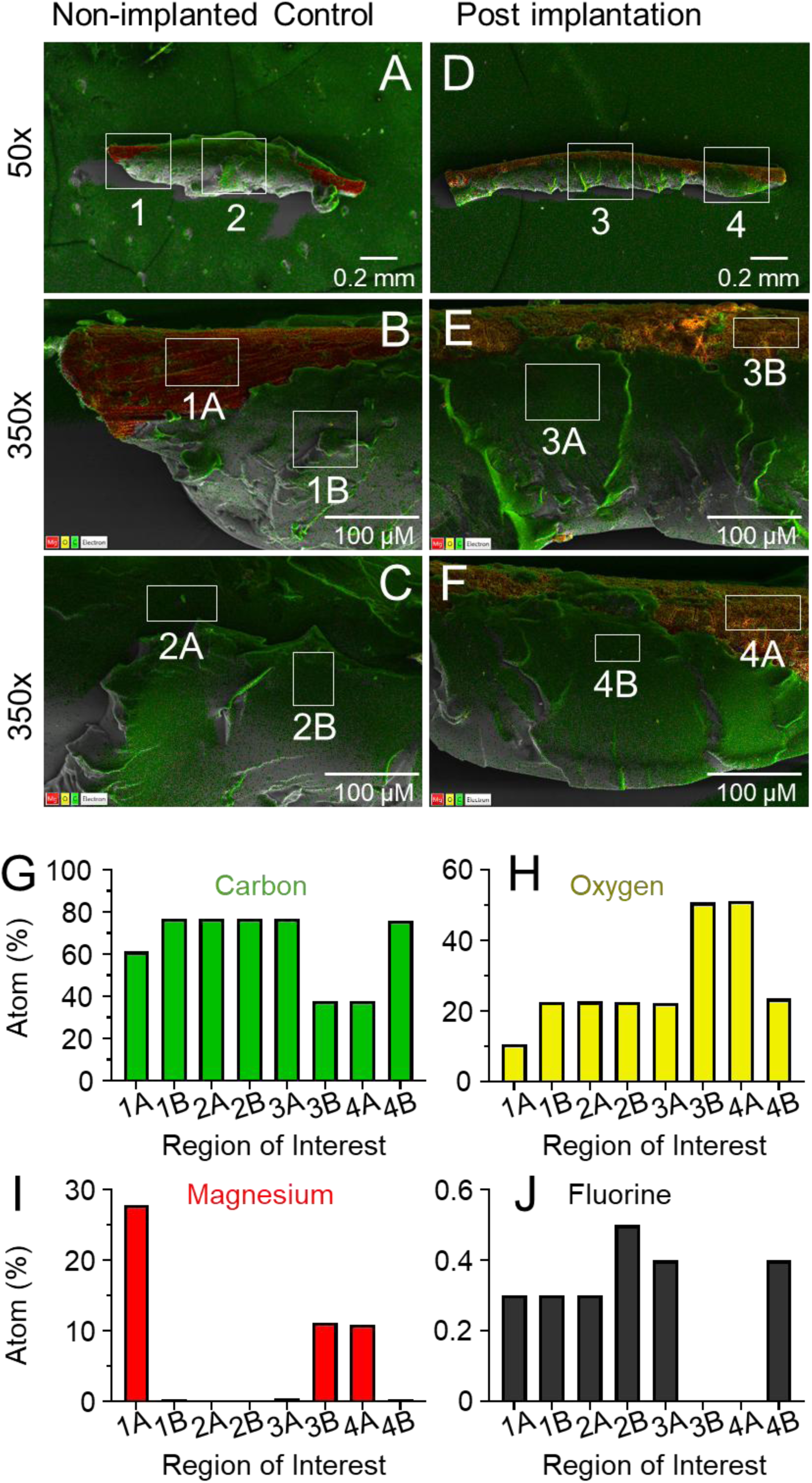
Representative SEM-EDS images of control and explanted devices. A,D) 50x gross SEM-EDS images of non-implanted control (A) vs explanted (D) IADD devices. The white squares indicate regions for higher magnification examination. B,C,E,F) 350x magnification SEM-EDS overlay images of non-implanted control (B,C) and explanted (E,F) devices. The regions of interest (white squares) indicate regions used to analyze the elemental distributions. G-J) Elemental distributions of carbon (G), oxygen (H), magnesium (I), and fluorine (J). The pseudo-colors in the SEM-EDS overlay (A-F) match the color conventions in (G-J). The orange color in (E,F) reflects the colocalization of magnesium (red) and oxygen (yellow).

## 4. Discussion

### 4.1 Design Considerations of the IADD Devices for Focal Drug Delivery

The present study demonstrates the loading of IADD devices with two model drugs, DEX and CIS, characterization of in vitro release and bioactivity, and IADD-mediated focal delivery of DEX to the kidney and brain. Selection of candidate drugs for IADD delivery should consider several factors. First, the amounts of drugs that can be loaded to the IADD devices are inherently limited by the sizes of arteries; thus, the selected therapeutic agents must have a therapeutic window at relatively low dosage levels, to achieve the intended clinical responses. Second, the risk-benefit balance is likely to be most favorable for drugs where focal delivery could meaningfully reduce systemic exposure or off-target toxicity. Third, the candidate therapeutic agents should be compatible with the device materials as well as the device fabrication and drug-loading processes Lastly, from an absorption, distribution, metabolism, and excretion (ADME) perspective, the candidate agents chosen for IADD should have pharmacologic activity in the target tissue without requiring hepatic conversion to an active metabolite and should have distribution properties that allow regional delivery to increase local exposure relative to systemic exposure. In this work, IADD devices consisting of bioresorbable Mg microwire substrates and biodegradable PGS polymer coatings were fabricated with helical or linear geometries, both of which supported proof-of-concept focal drug delivery in vivo. We believe that other biodegradable biocompatible metals (e.g., Zn, Mn, Mo, Fe, and their alloys) and other biodegradable generally-recognized-as-safe polyesters (or hydrogels) can be used to fabricate the IADD devices with other geometries (such as half-to-full ring or tubular structures or other stent-like mesh structures) to control the device degradation, modify drug loading, and achieve tunable drug release profiles for different therapeutic agents. However, the present study was not designed to isolate the individual effects of device geometry, loading method, and implantation site. Future studies should systematically vary these to define how IADD design influences release kinetics, focal biodistribution, vascular compatibility, and downstream tissue exposure.

In this work, the selected drugs (DEX and CIS) are compatible with the IADD device materials, i.e., magnesium (Mg) substrates (core backbone) and poly (glycerol sebacate) (PGS) coatings (drug-loading matrix), and processing parameters. Specifically, the drugs had at least moderate solubility in the solvents used for fabrication, such as ethanol or N, N-Dimethylformamide (DMF), to facilitate efficient loading without compromising the structural integrity of the device and pharmacological activity. Drug physicochemical properties are also expected to influence release behavior. Molecular weight, lipophilicity, charge, crystallinity, diffusivity through crosslinked PGS, and drug-polymer interactions may all affect burst release, sustained elution, and total recoverable payload. Accordingly, future studies should evaluate candidate drugs using matched device geometries and loading conditions, together with direct loading/recovery measurements and spatial drug-distribution analyses, to determine how drug properties influence IADD release kinetics.

### 4.2 Drug Loading, Release and Cellular Responses In Vitro

Elemental and thermal analyses supported drug incorporation and stability of both DEX and CIS within the PGS matrix. The observed cumulative release over 30 or 50 days shows that the polymer matrix can sustain extended elution while preserving the mechanical integrity of the Mg core for the duration of drug release from the IADD devices. In two-step DEX-loaded devices, DEX release was characterized by a pronounced day 1 burst followed by lower-level continued release over the remaining 30-day period. In contrast, CIS-loaded devices showed a more gradual release profile without a comparable burst phase. The one-step DEX-loaded devices showed quantifiable DEX release through day 50, with a lower relative day 1 burst than the two-step DEX-loaded devices. The distinct DEX and CIS release profiles suggest that drug release from IADD devices is governed by multiple interacting factors. The pronounced early DEX burst may reflect a higher fraction of drug localized near the PGS surface during post-soak loading, weaker retention within the crosslinked PGS matrix, and faster partitioning into the release medium. In contrast, the more gradual CIS release may result from stronger drug-matrix interactions, lower diffusivity through PGS, differences in molecular structure and ion coordination. Because DEX and CIS also differed in loading solvent, the present study cannot isolate the relative contribution of each factor. Recent molecular-scale analyses of PGS-based drug delivery similarly suggest that drug-polymer interactions, hydration, absorption, diffusion, and drug-specific physicochemical properties can influence release behavior. These studies support the need for future IADD optimization using drug-specific release testing, spatial drug-distribution analysis, and direct loading/recovery quantification [28]. Future studies will systematically vary drug physicochemical properties, loading solvent, PGS crosslinking density, and release medium, and will use spatial drug-mapping and direct loading quantification to clarify the mechanisms controlling burst release and sustained elution.

The absence of abrupt Mg²⁺ ion spikes throughout the 30-day incubation suggests that the PGS coating delayed rapid Mg degradation and helped preserve overall device integrity over the release period. The modest rise in Mg²⁺ concentration indicates that the protective effect of the PGS coating may decrease at later time points. This delayed increase in Mg²⁺ could result from progressive surface erosion of PGS, which increased water diffusion and ion release through the PGS layer under prolonged immersion. The two drug-loading methods used in this study may also contribute to differences in drug distribution, release kinetics, and in vivo targeting efficiency. In the two-step post-soaking method, drug loading likely occurs through solvent-mediated diffusion into the already crosslinked PGS matrix, which may enrich drug near the device surface and contribute to early burst release, particularly for DEX. In contrast, the one-step co-curing method disperses DEX into the pre-PGS before crosslinking, which may produce a more homogeneous drug distribution throughout the polymer matrix and reduce surface-localized drug accumulation. Potential advantages of the one-step method include more direct control over nominal drug loading and potentially more uniform drug distribution throughout the polymer. However, the two-step method may be more suitable for drugs that are unstable under the current curing conditions.

To evaluate cytotoxicity and released-drug bioactivity, two distinct in-vitro cell models were used. HUVECs represent the endothelial cells at the implantation site of IADD devices, while F98 glioma cells represent the cancer cells at the targeted organ site intended for focal drug treatment. In the HUVEC culture, cell viability remained above 90% for all device and control groups, supporting short-term endothelial cytocompatibility under the in vitro conditions tested. In contrast, in the F98 assay, the release media from the two-step CIS-loaded IADD device caused a 38% reduction in glioma cell viability, indicating inhibitory effect against cancer cells. The differences in the outcomes between HUVEC and F98 assays can be interpreted in the light of their respective *in vitro* culture methods that represent different in vitro exposure contexts. In the HUVEC assay, the cells were exposed to the intact device initially and then its dynamic degradation products in the same culture well gradually, where drug availability is governed by the slow degradation of the PGS matrix and localized diffusion from the device surface. This creates a concentration gradient, with HUVECs slowly getting exposed to CIS that is released into the immediate environment over 24 h. Thus, the amount of drug exposure at a given moment is well below the cytotoxic level, similar to the *in vivo* condition in an artery where continuous blood flow moderates the dynamic exposure level. By contrast, F98 glioma cells were exposed directly to the accumulated 24-hour release media, representing the total amount of CIS released from the IADD device in 24 hours. This results in higher effective drug concentrations than that in the exposure culture of HUVECs, and, thus, lowered cancer cell viability, similar to the expected higher drug levels at the targeted organ site *in vivo*. Overall, these *in vitro* results suggest the feasibility of IADD devices for focal drug delivery to a downstream organ.

### 4.3 Localized Focal Drug Delivery and in Vivo Performance in a Rat Model

Our *in vivo* results demonstrated the successful focal drug delivery to both kidney and brain - organs with distinct perfusion and barrier properties. For the smaller linear IADD devices, kidney and brain targeting achieved 109-fold and 68-fold (NODL) improvements in focal drug delivery, respectively. The brain targeting data demonstrates an intra-arterial concentration advantage even across a restrictive barrier. These outcomes align with pharmacokinetic theory for regional intra-arterial delivery, that is, high extraction/clearance favoring organ deposition. The localized gains observed in these studies can be leveraged in two complementary ways: (1) maintain standard target organ drug levels while substantially reducing systemic drug levels, and (2) substantially increase target organ drug levels while maintaining standard systemic drug levels. The optimal balance of each affordance will depend on the specific clinical target.

For intra-arterial delivery, local drug exposure depends on arterial input, downstream tissue extraction, tissue retention, regional blood flow, and systemic clearance. As described by Dedrick’s kinetic framework, regional delivery is expected to provide the greatest pharmacokinetic advantage when a drug exhibits favorable local extraction and systemic clearance properties [44]. Similarly, computational models of intra-arterial brain delivery emphasize that rapid tissue extraction and sustained retention can increase regional deposition [45]. In the present study, elevated DEX levels in the targeted kidney and brain relative to serum are consistent with these principles. And yet, the current single-timepoint biodistribution data do not define full pharmacokinetic profiles. Detectable DEX levels in the contralateral kidney and liver indicate that IADD-mediated delivery did not completely eliminate systemic redistribution. Future optimization of device drug loading, release kinetics, and drug formulation may help increase target-organ retention while reducing systemic leakage.

IADD-mediated focal drug delivery may be particularly advantageous for the disease processes that benefit from localized drug treatment, such as tumors, infection, inflammation, and pain. Previous studies have shown that intra-arterial delivery can restrict the drug distribution to the target region and thus reduce systemic toxicity [46, 47]. By enhancing the pharmacokinetic control of our IADD devices, we can leverage the advantages of IADD for sustained and precise therapeutic effects.

### 4.4 Limitations and Future Directions

Several limitations must be addressed to advance IADD devices toward translational development. The current drug-loading capacity is constrained by the solubility of therapeutic agents, the size of the target artery, and processing steps inherent to PGS as the polymer matrix. Thus, ideal candidate drugs should have high potency and low therapeutic dose requirements. In addition, the observed initial burst release of DEX indicates that further formulation optimization will be needed to achieve release profiles tailored to specific therapeutic applications. Although the linear IADD devices increased focal drug delivery to the target kidney and brain by 109-fold and 68-fold respectively, the DEX levels in the other compartments were still slightly elevated, indicating possible systemic leakage and associated off-target side effects. Lastly, vascularization of certain tissues, such as the cornea or striatum, poses challenges for the specificity of intra-arterial drug delivery to these areas, because there are no major feeding arteries specific to these tissues.

Although CIS-loaded IADD devices demonstrated in vitro release and retained bioactivity against glioma cells, CIS-IADD devices were not evaluated in vivo in the present study. Therefore, the current CIS data support in vitro feasibility but do not establish in vivo CIS biodistribution, antitumor efficacy, or toxicity. Future studies should evaluate CIS-IADD pharmacokinetics, target-organ biodistribution, tumor efficacy, and local/systemic safety in tumor-bearing models.

Intra-arterial implants carry established risks including thrombosis, embolism, and vessel occlusion. In our current study, histological examination of the implanted arteries revealed preservation of lumen and intact intima architecture, with no evidence of thrombus formation or luminal occlusion in the sections harvested at day 7. We additionally examined a small 55-day DEX-IADD implantation cohort, in which arterial histology showed preserved vessel morphology and the implanted device was no longer grossly evident in the examined arterial region. TTC staining of downstream brain tissue did not reveal gross infarct regions in the examined animals. Although the 7-day and 55-day arterial histology data provide preliminary evidence of local vascular tolerability, they do not fully assess all aspects of long-term vascular biocompatibility or systemic safety. Future safety studies should include larger cohorts, serial vascular histology, quantitative inflammatory and remodeling markers at the implantation site, hematology and serum chemistry analyses to assess systemic inflammation and organ toxicity, and measurements of Mg ion accumulation or distribution in major organs. Although the linear IADD device caused only limited cross-sectional obstruction, this estimate does not directly assess its effects on local hemodynamics. Device geometry may influence wall shear stress, flow separation, recirculation zones, and thrombosis risk. Therefore, future studies should incorporate computational fluid dynamics simulations and longitudinal in vivo vascular safety analyses to evaluate how IADD device shape and orientation affect local blood flow and vascular remodeling. The current study also did not directly correlate arterial histology with the corresponding explanted device surface. Although our data provide complementary evidence (Figs. 9 and 10), they do not define the spatial relationship between the device-contact region, residual PGS, local inflammatory or remodeling responses, and device erosion. Future studies should use in situ tissue sections containing the device or paired tissue-device surface analyses to more comprehensively evaluate the tissue–material interface.

A further limitation is that the current in vivo comparison was not dose-matched between IADD and oral DEX administration. Although NODL provides a useful route-normalized measure of organ-to-serum enrichment, it does not replace full pharmacokinetic analysis using matched doses and serial sampling. Future studies should include dose-normalized pharmacokinetic comparisons with oral and intravenous DEX administration, and potentially other focal delivery approaches, to more comprehensively define the pharmacokinetic advantage of IADD-mediated delivery. The current study also does not establish a direct in vitro-in vivo correlation because in vitro release was monitored for 30 or 50 days, whereas in vivo biodistribution was assessed only at day 7. Moreover, in vitro immersion conditions do not fully reproduce arterial blood flow, tissue uptake, metabolism, protein binding, or in vivo polymer degradation. Therefore, the 30-day release profiles should be interpreted as controlled in vitro performance data rather than direct predictors of in vivo pharmacokinetics. Future studies should include serial in vivo PK sampling, residual drug quantification in explanted devices, and flow-based in vitro release models to better correlate release kinetics with organ accumulation and systemic exposure. Recent endovascular drug-delivery and intravascular release-transport modeling studies further highlight the importance of coupling device release kinetics with blood flow, arterial-wall transport, and downstream tissue uptake when predicting local and systemic drug exposure [48, 49].

Future research should focus on enhancing drug loading efficiency and optimizing drug release profiles for different clinical applications. For example, tailoring polymer design by incorporating other polymers such as polycaprolactone (PCL) [50], polylactic acid (PLA) [51], poly(glycolic acid) (PGA), biodegradable hydrogels [52–55], silk, hybrid co-polymers, or their combinations could improve drug loading and release profiles. Moreover, incorporating 3D structural designs such as interconnected porous structures or meshes and multi-layer systems can also increase drug loading and sustain longer release. In addition, our current estimation methods for drug-loading (based on EDS atomic percentages and TGA residual mass) underestimate actual loaded mass, so development of more accurate loading quantification (e.g., direct extraction, imaging-based distribution mapping) is needed to better define therapeutic payloads. Additionally, combining PGS with angiogenic factors can improve tissue perfusion and enable effective drug delivery to poorly vascularized tissues. To reduce systemic leakage, optimizing drug formulations and delivery parameters are essential, potentially through advanced encapsulation techniques or refined device architectures.

Future studies will include dedicated histopathologic and functional evaluations in disease models, such as primary brain tumors. An important future direction is to assess organ-specific toxicities from our intra-arterial intervention. This will be particularly important in tumor models, which are less resilient to chemical and ischemic stressors [56]. By implementing these approaches, future iterations of IADD devices may achieve enhanced therapeutic efficacy, safety, and broad clinical applicability.

## 5. Conclusion

This study introduces biodegradable Mg/PGS IADD devices as a proof-of-concept approach for focal intra-arterial drug delivery. Prototype devices loaded with DEX or CIS demonstrated drug incorporation, in vitro release, endothelial cytocompatibility, and released-drug bioactivity. In vivo, DEX-loaded devices increased drug levels in downstream kidney and brain relative to systemic exposure, supporting focal target-organ enrichment. Additional studies will be required to define dose-normalized pharmacokinetics, optimize release profiles, evaluate long-term vascular and systemic safety, and determine therapeutic efficacy in disease models.

## Acknowledgements

This research project was supported by U.S. National Institutes of Health (NIBIB Grant EB035750) to EZ and HL. The authors appreciate the Central Facility for Advanced Microscopy and Microanalysis (CFAMM) for the use of SEM and EDS at the University of California at Riverside (UCR). We also thank the UC Riverside Metabolomics Core Facility at UCR for the Mass Spectrometry data acquisition support. The illustrations were created using BioRender (https://BioRender.com).

## Supplementary Methods

### Detailed fabrication of IADD devices with distinct shapes and loading methods

Two structural designs of IADD devices - helical and linear - were developed as first-generation prototypes to study drug release and biocompatibility in vitro and in vivo (Supplementary Figure 1). Initially, helical IADD devices were fabricated using a two-step post-soaking method, but their high release rate prompted the development of smaller, linear configurations with refined loading strategies. Accordingly, the helical two-step devices were used for renal artery implantation (kidney cohort 1), the linear two-step devices were used for brain delivery and in vitro studies, and the linear one-step devices were used for renal artery delivery (kidney cohort 2).

Magnesium wire (Sigma-Aldrich, 127 µm diameter) was cut into 4 mm segments and ultrasonically cleaned sequentially in acetone and ethanol. PGS pre-polymer was synthesized using a modified procedure from Wang et al., 2002. Briefly, 60.67 g sebacic acid (Thermo Scientific, 98%) and 27.62 g glycerol (Fisher Chemical, 99.5%) were heated at 120 °C under nitrogen for 2 h to melt the sebacic acid, followed by polymerization at 120 °C and 300 rpm for 24 h under nitrogen, and then 42 h under vacuum to yield pre-PGS. The cooled product (70 °C) was poured into chilled (4 °C) Millipore water to remove residual glycerol, washed three times, freeze-dried at -40 °C for 48 h, and stored at -20 °C. Viscosity was measured with a rheometer (Anton Paar MCR92) between 0-120 °C.

For linear devices, approximately 2 g of pre-PGS was poured into an 82 × 36 mm Teflon mold (∼600 µm layer), and cleaned Mg segments were embedded before vacuum curing at 120 °C for 120 h (Across International AT32). The cured composite was cut under a stereomicroscope into 400 × 250 × 4200 µm devices, maintaining a central Mg core.

In the two-step post-soaking method, the cured devices were soaked in 100% ethanol for 24 h to remove residual sebacic acid, then immersed for 48 h in either DEX (10 mg/mL in ethanol) or CIS (15 mg/mL in DMF) solutions. Devices soaked in ethanol or DMF without drug served as IADD controls. All devices were vacuum-dried at 50 °C for 2 h and UV-sterilized for 30 min.

Helical IADDs were fabricated to provide increased surface area and drug-loading capacity. Mg wire (127 µm × 25 mm) was wound around a 650 µm stainless-steel mandrel to form a 3.8 mm-long helix and placed into a 1 mm-diameter Teflon mold. A 400 µm stainless-steel rod was inserted centrally to preserve a hollow lumen for arterial blood flow. It was then cured under vacuum at 120 °C for 120 h, and the helical devices were trimmed to 4 mm length. These were loaded using the 2-step post-soaking method described above.

In the one-step co-curing method, DEX (10% w/w) was directly dispersed into pre-PGS at 70 °C under stirring in a speed mixer (3000 rpm), followed by insertion of the Mg wire and vacuum curing at 120 °C for 120 h. This process achieved uniform drug distribution throughout the PGS matrix and eliminated the need for post-fabrication soaking.

### Detailed methodology for measuring DEX and CIS levels using LC/MS-MS

To measure the DEX and CIS released into the media, liquid chromatography tandem mass spectrometry (LC-MS/MS) was used. For DEX estimation, 200 µL of ethyl acetate and 25 ng/ml of dexamethasone-d5 (internal standard) were added to 50 µL of salt solutions. The mixture was vortexed, centrifuged (14000 rpm, 4 °C, 10 min), dried under air, and resuspended in 200 µL of methanol. The samples were analyzed using a 50 mm Acquity BEH C-18 column on a Waters I-class UPLC coupled to a Synapt G2-Si Q-TOF mass spectrometer (Waters Corporation, Milford, MA, USA). The mobile phases were (A) water with 0.1% formic acid and (B) acetonitrile with 0.1% formic acid. The flow rate was 400 µL/min, and the column was held at 40 °C. The data acquisition utilized an injection volume of 5 µL. The gradient was as follows: 0 min, 0% B; 1.5 min, 0% B; 2.5 min, 95% B; 4.5 min, 95% B; 4.75 min, 0% B; 6.25 min, 0% B. Target compound responses were normalized against the deuterated internal standard, and the amounts were calculated using a standard curve. The LC-MS data were acquired in positive electrospray ionization (ESI+) mode for a mass range from m/z 50 to 1000 with a scan time of 0.05 s. A lockspray reference ion with m/z 556.2771, corresponding to 1 ng/µL leucine enkephalin, was infused at a rate of 5 µL/min to achieve real-time mass correction during the data acquisition. The ESI mass spectra were acquired in positive mode using a capillary voltage of 1.5 kV and source temperature of 150 °C. Desolvation was achieved by directing nitrogen at 600 °C, 1000 L/h. After evaluating the full scan TOF MS data for the targeted ions (m/z 398.2379, m/z 473.1735, and m/z 393.2037), three distinct TOF MS/MS channels were developed in the method. These channels were utilized for identification and data analysis using unique fragment ions.

The CIS in the salt solutions was analyzed as described previously (Shaik et al., 2017). Briefly, 5 µl of palladium acetate was added to 50 µl of collected physiological salt solution (PSS; rSBF or aCSF) followed by 15 µl of 1% v/v diethyldithiocarbamate (DTCC) in 0.1N NaOH solution. The mixture was vortexed followed by incubation at 40 °C for 30 min in a water bath. Following incubation, 1.5 ml of acetonitrile containing 1 ng/ml of 8-Cyclopentyl-1,3-dipropylxanthine (DPCPX) (internal standard) was added to each tube. The samples were vortexed for 10 min and then centrifuged at 4000 rpm and the contents were dried under air. The dried samples were reconstituted with 80:20:0.1% water: acetonitrile: formic acid and loaded onto a Waters H-class UPLC coupled to G2-XS Q-TOF mass spectrometer (Waters Corporation, Milford, MA, USA) for the analysis. The mobile phases were (A) water with 0.1% formic acid and (B) acetonitrile with 0.1% formic acid. The flow rate was 350 µL/min, and the column was held at 25 °C. The data acquisition utilized an injection volume of 1 µL. The gradient was as follows: 0 min, 1% B, 0.5 min, 1% B, 1 min, 75% B, 1.5 min, 90% B, 2.5 min, 99% B, 6 min, 99% B, 8 min, 1% B and 10 min, 1% B. The samples were normalized against the internal standard, and the amount of CIS was calculated using the standard curve. The LC-MS data were acquired in positive electrospray ionization (ESI+) mode for a mass range from m/z 50 to 800 with a scan time of 0.3 s. A lockspray reference ion with m/z 556.2771, corresponding to 1 ng/µL leucine enkephalin, was infused at a rate of 10 µL/min to achieve real-time mass correction during the data acquisition. The ESI mass spectra were obtained in positive mode with a capillary voltage of 1.0 kV, a source temperature of 150 °C, and a cone gas (N2) flow rate of 50 L/h. Desolvation was achieved by directing nitrogen at 600 °C, 850 L/h. After evaluating the full scan ToF MS data for the targeted ions (m/z 492.014, m/z 305.198, and m/z 402.966), three distinct ToF MS/MS channels were developed in the method. These channels were utilized for identification and data analysis using unique fragment ions. Data were processed using the Waters Quanlynx application.

---

**Supplementary Figure S1.**
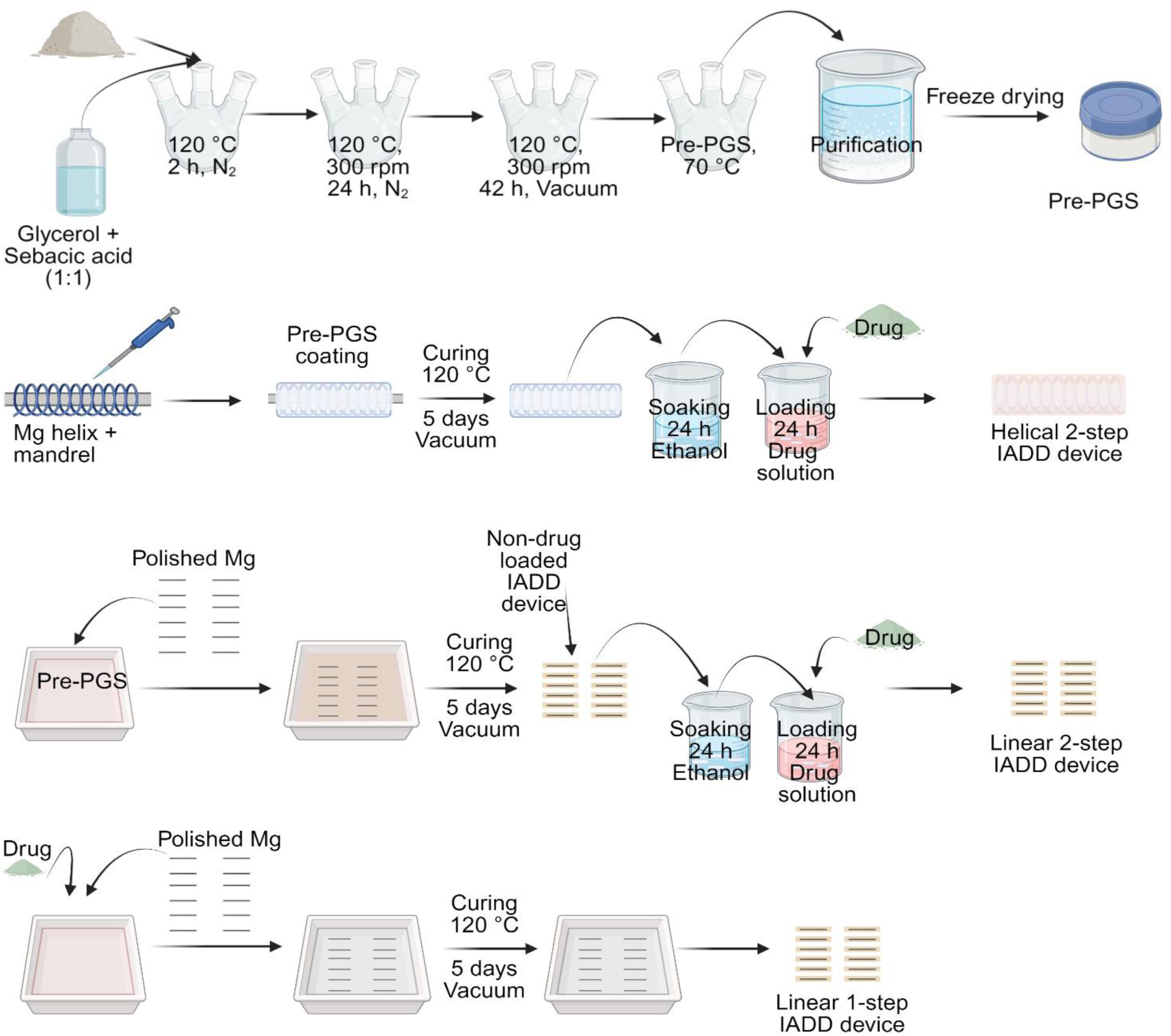
Different schemes for IADD device fabrication.

**Supplementary Figure S2.**
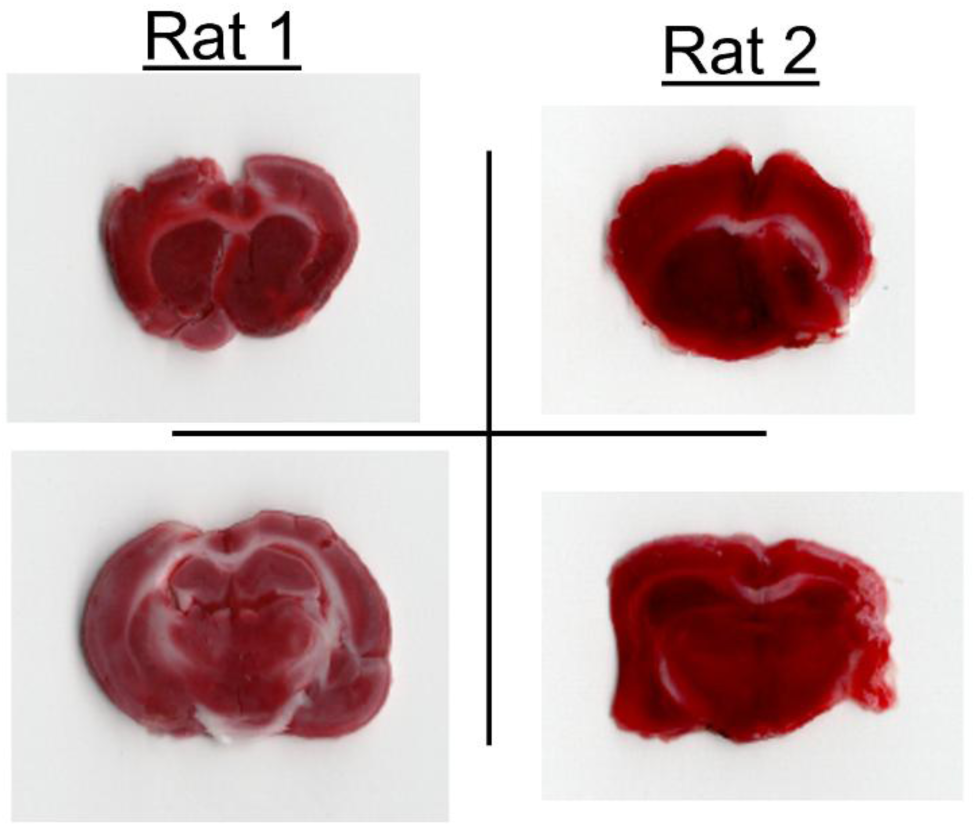
Gross assessment of ischemic injury of the brain by TTC staining after DEX-IADD implantation. Representative TTC-stained coronal brain sections from two rats collected at the 55-day endpoint. Brain sections showed overall preserved TTC staining without obvious macroscopic infarct-like pallor in the examined sections.

**Supplementary Table 1.** Comparison of in vitro release profiles for one-step and two-step IADD devices. DEX-IADD and CIS-IADD devices were compared based on day 1 release, cumulative release through day 7, cumulative release total, the corresponding percentages of total recovered drug, and mean daily release after day 7. Values are reported as mean ± SD.

| Device | Loading method | Drug | Day 1 release ($\mu\text{g}$) | Cumulative release through day 7 ($\mu\text{g}$) | Cumulative release total ($\mu\text{g}$) | Day 1 release (% of total recovered drug) | Day 1-7 release (% of total recovered drug) | Mean daily release after day 7 ($\mu\text{g}/\text{day}$) |
| --- | --- | --- | --- | --- | --- | --- | --- | --- |
| DEX-IADD | Two-step | DEX | 274.65 ± 91.90 | 331.95 ± 98.93 | 373.11 ± 98.20 | 73.67 ± 4.42 | 88.84 ± 2.41 | 1.77 ± 0.14 |
| DEX-IADD | One-step | DEX | 41.03 ± 9.70 | 120.04 ± 29.16 | 150.06 ± 34.97 | 27.92 ± 5.93 | 79.91 ± 3.28 | 0.99 ± 0.08 |
| CIS-IADD | Two-step | CIS | 4.49 ± 3.55 | 24.98 ± 5.99 | 64.73 ± 7.82 | 7.6 ± 4.93 | 35.53 ± 10.42 | 1.68 ± 0.14 |

## Bibliography

[1] R. Huang, J. Boltze, S. Li, Strategies for improved intra-arterial treatments targeting brain tumors: a systematic review, Frontiers in Oncology, 10 (2020) 1443.

[2] J.A. Ellis, M. Banu, S.S. Hossain, R. Singh-Moon, S.D. Lavine, J.N. Bruce, S. Joshi, Reassessing the role of intra-arterial drug delivery for glioblastoma multiforme treatment, Journal of drug delivery, 2015 (2015) 405735.

[3] J.S. Speed, K.A. Hyndman, In vivo organ specific drug delivery with implantable peristaltic pumps, Scientific reports, 6 (2016) 26251.

[4] H.D. Jackson, M.J. Cotler, G.W. Saunders, C.A. Cornelssen, P.J. West, C.S. Metcalf, K.S. Wilcox, M.J. Cima, Intracerebral delivery of antiseizure medications by microinvasive neural implants, Brain, (2024) awae282.

[5] H.B. Newton, M.A. Page, L. Junck, H.S. Greenberg, Intra-arterial cisplatin for the treatment of malignant gliomas, Journal of neuro-oncology, 7 (1989) 39–45.

[6] E. Galanis, J. Buckner, Chemotherapy for high-grade gliomas, British journal of cancer, 82 (2000) 1371–1380.

[7] A.B. Crocker, M.H. Talukder, M.S. Ali, A.S. Calvino, P. Somasundar, N.J. Espat, Regional Hepatic Therapies for Colorectal Hepatic Metastases, Rhode Island Medical Journal, 108 (2025).

[8] S. Fu, A. Naing, S.L. Moulder, K.S. Culotta, D.C. Madoff, C.S. Ng, T.L. Madden, G.S. Falchook, D.S. Hong, R. Kurzrock, Phase I trial of hepatic arterial infusion of nanoparticle albumin–bound paclitaxel: toxicity, pharmacokinetics, and activity, Molecular cancer therapeutics, 10 (2011) 1300–1307.

[9] C. Ferreira, M.Y. Ferreira, F. Singh, T. Wong, S. Bokil, S. Massimo, J. Cavallaro, O. Albers, R. D’Amico, D. Langer, Superselective intra-arterial cerebral infusion of chemotherapeutics after osmotic blood–brain barrier disruption in newly diagnosed or recurrent glioblastoma: technical insights and clinical outcomes from a single-center experience, Journal of NeuroInterventional Surgery, (2025).

[10] S. Joshi, C.W. Emala, J. Pile-Spellman, Intra-arterial drug delivery: a concise review, Journal of Neurosurgical Anesthesiology, 19 (2007) 111–119.

[11] M. Pogorielov, E. Husak, A. Solodivnik, S. Zhdanov, Magnesium-based biodegradable alloys: Degradation, application, and alloying elements, Interventional Medicine and Applied Science, 9 (2017) 27–38.

[12] Z.-J. Sun, C. Chen, M.-Z. Sun, C.-H. Ai, X.-L. Lu, Y.-F. Zheng, B.-F. Yang, D.-L. Dong, The application of poly (glycerol–sebacate) as biodegradable drug carrier, Biomaterials, 30 (2009) 5209–5214.

[13] C. Zhang, T.H. Wen, K.A. Razak, J. Lin, C. Xu, C. Seo, E. Villafana, H. Jimenez, H. Liu, Magnesium-based biodegradable microelectrodes for neural recording, Materials Science and Engineering: C, 110 (2020) 110614.

[14] R. Sheng, J. Mu, R.V. Chernozem, Y.R. Mukhortova, M.A. Surmeneva, I.O. Pariy, T. Ludwig, S. Mathur, C. Xu, R.A. Surmenev, Fabrication and characterization of piezoelectric polymer composites and cytocompatibility with mesenchymal stem cells, ACS Applied Materials & Interfaces, 15 (2023) 3731–3743.

[15] D. Tie, N. Hort, M. Chen, R. Guan, S. Ulasevich, E.V. Skorb, D. Zhao, Y. Liu, P. Holt-Torres, H. Liu, In vivo urinary compatibility of Mg-Sr-Ag alloy in swine model, Bioactive Materials, 7 (2022) 254–262.

[16] D. Tie, H. Liu, R. Guan, P. Holt-Torres, Y. Liu, Y. Wang, N. Hort, In vivo assessment of biodegradable magnesium alloy ureteral stents in a pig model, Acta Biomaterialia, 116 (2020) 415–425.

[17] F. Witte, The history of biodegradable magnesium implants: a review, Acta biomaterialia, 6 (2010) 1680–1692.

[18] N. Banjanin, G. Belojevic, Changes of blood pressure and hemodynamic parameters after oral magnesium supplementation in patients with essential hypertension—an intervention study, Nutrients, 10 (2018) 581.

[19] A. Yamamoto, S. Hiromoto, Effect of inorganic salts, amino acids and proteins on the degradation of pure magnesium in vitro, Materials Science and Engineering: C, 29 (2009) 1559–1568.

[20] I. Johnson, H. Liu, A study on factors affecting the degradation of magnesium and a magnesium-yttrium alloy for biomedical applications, PLoS One, 8 (2013) e65603.

[21] Y. Wang, G.A. Ameer, B.J. Sheppard, R. Langer, A tough biodegradable elastomer, Nature biotechnology, 20 (2002) 602–606.

[22] C.G. Hynes, E. Morra, P. Walsh, F. Buchanan, Degradation of biomaterials, in: Tissue Engineering, Elsevier, 2023, pp. 213–259.

[23] I. Pomerantseva, N. Krebs, A. Hart, C.M. Neville, A.Y. Huang, C.A. Sundback, Degradation behavior of poly (glycerol sebacate), Journal of Biomedical Materials Research Part A: An Official Journal of The Society for Biomaterials, The Japanese Society for Biomaterials, and The Australian Society for Biomaterials and the Korean Society for Biomaterials, 91 (2009) 1038–1047.

[24] B. Godinho, N. Gama, A. Ferreira, Different methods of synthesizing poly (glycerol sebacate)(PGS): A review, Frontiers in Bioengineering and Biotechnology, 10 (2022) 1033827.

[25] R. Martín-Cabezuelo, J.C. Rodríguez-Hernández, G. Vilariño-Feltrer, A. Vallés-Lluch, Role of curing temperature of poly (glycerol sebacate) Substrates on protein-cell interaction and early cell adhesion, Polymers, 13 (2021) 382.

[26] S. Andrä-Żmuda, P. Chaber, M. Martinka Maksymiak, M. Musioł, G.Y. Adamus, Poly (glycerol sebacate): A Comparative Study of Various Synthesis Methods, Biomacromolecules, (2025).

[27] M.-L.L. Bice, M.H. Yu, V.L. Ortega, C.-C. Hsu, K.J. McHugh, Methacrylated poly (glycerol sebacate) as a photocurable, biocompatible, and biodegradable polymer with tunable degradation and drug release kinetics, Drug delivery and translational research, 15 (2025) 2694–2709.

[28] X. Davoy, J. Devémy, P. Fayon, P. Chennell, M. Sahihi, S. Garruchet, A. Dequidt, P. Hauret, P. Malfreyt, Drug Delivery Mechanisms of Poly (glycerol sebacate): An In-Depth Study of the Energetics at the Molecular Scale, Molecular Pharmaceutics, 22 (2025) 3848–3859.

[29] S. Salehi, M. Czugala, P. Stafiej, M. Fathi, T. Bahners, J.S. Gutmann, B.B. Singer, T.A. Fuchsluger, Poly (glycerol sebacate)-poly (epsilon-caprolactone) blend nanofibrous scaffold as intrinsic bio- and immunocompatible system for corneal repair, Acta Biomater, 50 (2017) 370–380.

[30] B.M. Hall, Corticosteroids in autoimmune diseases, Australian Prescriber, 22 (1999) 9.

[31] P. Barnes, Molecular mechanisms of corticosteroids in allergic diseases, Allergy, 56 (2001).

[32] I.A. Janahi, A. Rehman, N.U.-A. Baloch, Corticosteroids and their use in respiratory disorders, Corticosteroids; InTech Open: London, UK, (2018) 47–57.

[33] R.S. Goodman, D.B. Johnson, J.M. Balko, Corticosteroids and cancer immunotherapy, Clinical Cancer Research, 29 (2023) 2580–2587.

[34] V.S. Madamsetty, R. Mohammadinejad, I. Uzieliene, N. Nabavi, A. Dehshahri, J. Garcia-Couce, S. Tavakol, S. Moghassemi, A. Dadashzadeh, P. Makvandi, A. Pardakhty, A. Aghaei Afshar, A. Seyfoddin, Dexamethasone: Insights into Pharmacological Aspects, Therapeutic Mechanisms, and Delivery Systems, ACS Biomater Sci Eng, 8 (2022) 1763–1790.

[35] M.E. Cooley, L. Davis, J. Abrahm, Cisplatin: a clinical review. Part II--Nursing assessment and management of side effects of cisplatin, Cancer Nurs, 17 (1994) 283–293.

[36] J.H. An, Y. Su, T. Radman, M. Bikson, Effects of glucose and glutamine concentration in the formulation of the artificial cerebrospinal fluid (ACSF), Brain research, 1218(2008) 77–86.

[37] A. Oyane, H.M. Kim, T. Furuya, T. Kokubo, T. Miyazaki, T. Nakamura, Preparation and assessment of revised simulated body fluids, Journal of Biomedical Materials Research Part A: An Official Journal of The Society for Biomaterials, The Japanese Society for Biomaterials, and The Australian Society for Biomaterials and the Korean Society for Biomaterials, 65 (2003) 188–195.

[38] C. Xu, Y. Chen, J. Lin, H.H. Liu, Direct and indirect culture methods for studying biodegradable implant materials In Vitro, Journal of Visualized Experiments (JoVE), (2022) e63065.

[39] V. Vichai, K. Kirtikara, Sulforhodamine B colorimetric assay for cytotoxicity screening, Nat Protoc, 1 (2006) 1112–1116.

[40] A.T. Feldman, D. Wolfe, Tissue processing and hematoxylin and eosin staining, in: Histopathology: methods and protocols, Springer, 2014, pp. 31–43.

[41] S. Oh, X.L. Mai, J. Kim, A.C.V. de Guzman, J.Y. Lee, S. Park, Glycerol 3-phosphate dehydrogenases (1 and 2) in cancer and other diseases, Experimental & molecular medicine, 56 (2024) 1066–1079.

[42] S.M. Madiraju, E. Possik, F. Al-Mulla, C.J. Nolan, M. Prentki, Glycerol and glycerol-3-phosphate: multifaceted metabolites in metabolism, cancer and other diseases, Endocrine Reviews, (2025) bnaf033.

[43] M.H. Andrews, S.A. Wood, R.J. Windle, S.L. Lightman, C.D. Ingram, Acute glucocorticoid administration rapidly suppresses basal and stress-induced hypothalamo-pituitary-adrenal axis activity, Endocrinology, 153 (2012) 200–211.

[44] A. Karssen, O. Meijer, A. Berry, R. Sanjuan Pinol, E. De Kloet, Low doses of dexamethasone can produce a hypocorticosteroid state in the brain, Endocrinology, 146 (2005) 5587–5595.

[45] L. Yang, K. Boyd, S.C. Kaste, L. Kamdem Kamdem, R.J. Rahija, M.V. Relling, A mouse model for glucocorticoid-induced osteonecrosis: effect of a steroid holiday, Journal of Orthopaedic Research, 27 (2009) 169–175.

[46] S. Çitoğlu, H. Duran, Recent advances in porous nanomaterials-based drug delivery systems for osteoarthritis, Nano Select, (2023).

[47] Z. Lu, L. Xie, W. Liu, Z. Li, Y. Chen, G. Yu, B. Shi, A bibliometric analysis of intra-articular injection therapy for knee osteoarthritis from 2012 to 2022, Medicine, 102 (2023) e36105.

[48] X. Feng, T. Jiang, Mathematical and numerical analysis for PDE systems modeling intravascular drug release from arterial stents and transport in arterial tissue, arXiv preprint arXiv:2404.13780, (2024).

[49] C.N. Lungu, A. Creteanu, M.C. Mehedinti, Endovascular drug delivery, Life, 14 (2024) 451.

[50] X. Yang, J. Wei, D. Lei, Y. Liu, W. Wu, Appropriate density of PCL nano-fiber sheath promoted muscular remodeling of PGS/PCL grafts in arterial circulation, Biomaterials, 88 (2016) 34–47.

[51] M. Frydrych, S. Román, S. MacNeil, B. Chen, Biomimetic poly (glycerol sebacate)/poly (l-lactic acid) blend scaffolds for adipose tissue engineering, Acta Biomaterialia, 18 (2015) 40–49.

[52] R. Daya, C. Xu, N.T. Nguyen, H.H. Liu, Angiogenic Hyaluronic Acid Hydrogels with Curcumin-Coated Magnetic Nanoparticles for Tissue Repair, ACS Appl Mater Interfaces, 14 (2022) 11051–11067.

[53] R. Raho, N.Y. Nguyen, N. Zhang, W. Jiang, A. Sannino, H. Liu, M. Pollini, F. Paladini, Photo-assisted green synthesis of silver doped silk fibroin/carboxymethyl cellulose nanocomposite hydrogels for biomedical applications, Mater Sci Eng C Mater Biol Appl, 107 (2020) 110219.

[54] C. Xu, C. Hung, Y. Cao, H.H. Liu, Tunable Crosslinking, Reversible Phase Transition, and 3D Printing of Hyaluronic Acid Hydrogels via Dynamic Coordination of Innate Carboxyl Groups and Metallic Ions, ACS Appl Bio Mater, 4 (2021) 2408–2428.

[55] N. Zhang, J. Lock, A. Sallee, H. Liu, Magnetic Nanocomposite Hydrogel for Potential Cartilage Tissue Engineering: Synthesis, Characterization, and Cytocompatibility with Bone Marrow Derived Mesenchymal Stem Cells, ACS Appl Mater Interfaces, 7 (2015) 20987–20998.

[56] T. Bae, S.P. Hallis, M.-K. Kwak, Hypoxia, oxidative stress, and the interplay of HIFs and NRF2 signaling in cancer, Experimental & molecular medicine, 56 (2024) 501–514.

